# Bacteriocin production in *Streptococcus gallolyticus* is controlled by a complex 4-component regulatory system with activator and anti-activator activities

**DOI:** 10.1101/2020.06.04.131722

**Authors:** Alexis Proutière, Bruno Périchon, Laurence du Merle, Hugo Varet, Patrick Trieu-Cuot, Shaynoor Dramsi

**Affiliations:** Unité de Biologie des Bactéries Pathogènes à Gram-positif, Institut Pasteur, Paris, France; CNRS Unité Mixte de Recherches, UMR 2001, Paris, France; Université Paris Diderot, Sorbonne Paris Cité, Paris, France; Institut Pasteur, Hub Bioinformatique et Biostatistique, Département de Biologie Computationnelle (USR 3756 IP CNRS), Paris, France; Institut Pasteur, Biomics Platform, Centre de Ressources et Recherches Technologiques (C2RT), Paris, France

## Abstract

Bacteriocins are natural antimicrobial peptides produced by bacteria to kill closely related competitors. The opportunistic pathogen *Streptococcus gallolyticus* (*Sgg*) was recently shown to outcompete commensal enterococci of the murine microbiota in tumoral conditions thanks to the production of a two-peptide bacteriocin named gallocin. We here identified 4 genes involved in the regulatory control of gallocin in *Sgg* UCN34, respectively encoding a histidine kinase/response regulator two-component system (BlpH/BlpR), a secreted peptide (GSP), and a putative regulator of unknown function (BlpS). While BlpR is a typical 243-aa response regulator possessing a phospho-receiver domain and a LytTR DNA-binding domain, BlpS is a 108-aa protein containing only a LytTR domain. Our results showed that the secreted peptide GSP activates the dedicated two-component system BlpH/BlpR to induce gallocin transcription. A genome-wide transcriptome analysis indicates that this regulatory system (GSP-BlpH/BlpR) is highly specific for bacteriocin production. Importantly, as opposed to BlpR, BlpS was shown to repress gallocin gene transcription. A conserved operator DNA sequence of 30-bp was found in all promoter regions regulated by BlpR and BlpS. EMSA assays showed direct and specific binding of the two gallocin regulators to various regulated promoter regions in a dose dependent manner. Gallocin expression appears tightly controlled in *Sgg* by quorum sensing and antagonistic activity of 2 LytTR-containing proteins.

**Significance:** *Streptococcus gallolyticus* (*Sgg*), formely known as *S. bovis* biotype I, is an opportunistic pathogen causing septicemia and endocarditis in the elderly often associated with asymptomatic colonic neoplasia. We previously showed that *Sgg* produces a bacteriocin, termed gallocin, enabling colonization of the colon in tumoral conditions by outcompeting commensal members of the gut. Here we characterized a 4-component regulatory system that regulates gallocin transcription, which is activated by the response regulator BlpR. BlpR itself is activated by a quorum sensing peptide GSP and a dedicated histidine kinase BlpH. Interestingly, BlpS, a small DNA-binding protein co-transcribed with BlpR was found to repress gallocin genes transcription, likely by antagonizing BlpR. Understanding gallocin regulation is crucial to prevent *Sgg* colon colonization in tumoral conditions.

## Introduction

*Streptococcus gallolyticus* subspecies *gallolyticus* (*Sgg*), formerly known as *Streptococcus bovis* biotype I, is an opportunistic Gram-positive pathogen responsible for septicemia and endocarditis in the elderly (1). Invasive *Sgg* infections are strongly associated with asymptomatic colonic neoplasia but the mechanisms underlying this association are still unclear (2, 3). Recently, it was shown that *Sgg* produces a specific bacteriocin named gallocin, whose antimicrobial activity is enhanced by the increased level of secondary bile salts observed in tumoral conditions, allowing *Sgg* to colonize the murine gut by killing resident enterococci (4). As such, gallocin constitutes the first bacterial factor which could explain the *Sgg* association with colonic tumors.

Bacteriocins are natural antimicrobial peptides produced by many bacteria. Producer strains are protected from their own bacteriocin by the presence of an immunity system. Most bacteriocins have a narrow spectrum of activity restricted to bacteria closely related to the producer. Therefore, bacteriocin production is important for the colonization of specific niches, especially in competitive environments such as the gut (5). Bacteriocins of gram-positive bacteria have been divided into four classes based on size, amino acid composition, and structure (6). Class I includes small (< 5 kDa) linear peptides containing post-translationally modified amino acids called lantibiotics, class II small (< 10 kDa) linear peptides without post-translationally modified amino acids, class III large (> 10 kDa) proteins and class IV small cyclic peptides. Class II bacteriocins are further subdivided into three groups: class IIa consists of pediocin-like bacteriocins, class IIb of two or more peptides and class IIc of all other bacteriocins not fitting in IIa and IIb. *In silico* analysis indicates that gallocin likely belongs to class IIb bacteriocins (4). In general, these bacteriocins kill susceptible strains by forming pores in the target membranes, resulting in ion leakage and cell death (7).

Some class IIb bacteriocin loci encode a 3-component regulatory system composed of an inducing peptide and a dedicated two-component system (TCS) with a membrane-bound histidine kinase and a cytoplasmic response regulator. Activation of bacteriocin production through this regulatory system is similar to quorum-sensing regulatory systems. First, the inducing peptide is secreted into the extracellular medium and, upon reaching a threshold concentration, binds to and activates the histidine kinase, resulting in phosphorylation of its associated response regulator. The phosphorylated response regulator then activates the transcription of genes necessary for class IIb bacteriocin production, including its own transcription, resulting in a rapid overexpression of the regulated genes (7–9). In streptococci, complex regulatory cross-talk has been identified between bacteriocin production and competence (10). Natural competence has been reported in the related *S. bovis* (11) but not in *Sgg*. A peptide previously identified in the extracellular medium of *Sgg* called CSP (due to its similarity with Competence Stimulating Peptide) was shown to induce bacteriocin production but did not allow capture and integration of foreign plasmid DNA (12). In the accompanying paper, we demonstrate that CSP should be renamed GSP for Gallocin Stimulating Peptide.

The aim of the present study was to identify and characterize the regulatory system involved in gallocin production and to identify other potentially co-regulated genes. In addition to the three-component system, consisting of an inducing peptide, a histidine kinase and a response regulator, a fourth regulatory component allowing tight control of gallocin expression was identified in this work. Moreover, the presence of several putative novel bacteriocins co-expressed with gallocin highlights the importance of these antimicrobial peptides for the gut colonization by this pathobiont associated with colorectal cancer.

## Results

### 1 Identification of a dedicated three-component regulatory system involved in gallocin production

To understand how gallocin production is regulated in *Sgg* UCN34 (13), the genomic locus encoding this putative class IIb bacteriocin was inspected for the presence of potential regulatory genes. Genes encoding a 3-component regulatory system were identified at one end of the gallocin locus **(Fig. 1A)**. This module is composed of 3 genes: *blpH*, a putative histidine kinase, *blpR*, a putative response regulator and a divergently transcribed gene encoding a putative inducing peptide named *gsp* (gallocin-stimulating peptide). The regulatory genes are close to the genes encoding the gallocin peptides (*blpA* and *blpB*) and the putative immunity protein (*blpC*), two genes encoding an ABC transporter (*blpT1* and *blpT2*) shown to be involved in gallocin peptide secretion (Harrington-Proutière et al., 2020, companion paper), and other conserved bacteriocin-associated proteins such as Abi domain proteins (*gallo_rs10400-10405*) (**Fig. 1A)**. A genetic approach was undertaken to demonstrate the role of these three regulatory genes in gallocin production. Markerless in-frame deletion mutants of *gsp*, *blpH* and *blpR* were obtained in *Sgg* UCN34. For each mutant, we also selected a clone that reverted to the wild-type genotype (bWT) following homologous recombination. Gallocin production is easily visualized through its antimicrobial activity against the very closely related bacterium *Streptococcus gallolyticus* subsp. *macedonicus (Sgm*) (Sup. **Fig. S1A**), which was used as a susceptible indicator strain throughout this manuscript. As shown in **Fig. 1B**, gallocin production was abolished in the Δ*gsp*, Δ*blpH* and Δ*blpR* mutants as compared to their bWT strains. All three mutants did not exhibit any killing activity against the *Sgm* prey strain, indicating that these three genes are essential for gallocin production in *Sgg* UCN34. We reasoned that if *gsp* encodes a secreted inducing peptide that activates its cognate two-component system, addition of GSP peptide to the extracellular medium should restore gallocin production by the Δ*gsp* mutant. We also hypothesized that GSP, like other inducing peptides, is synthesized as a precursor matured by cleavage upon secretion after a double glycine motif (14). The predicted mature GSP peptide corresponding to the 24 C-terminal amino acids encoded by *gsp* was synthesized (**Fig. S1B**). Addition of synthetic GSP to the culture medium restored gallocin production by the Δ*gsp* mutant (**Fig. 1B**). Importantly, addition of GSP did not restored gallocin production in the Δ*blpH* or Δ*blpR* mutants, an epistatic relationship suggesting that GSP activates transcription of genes involved in gallocin production through the BlpH/R TCS.

**Figure 1:**
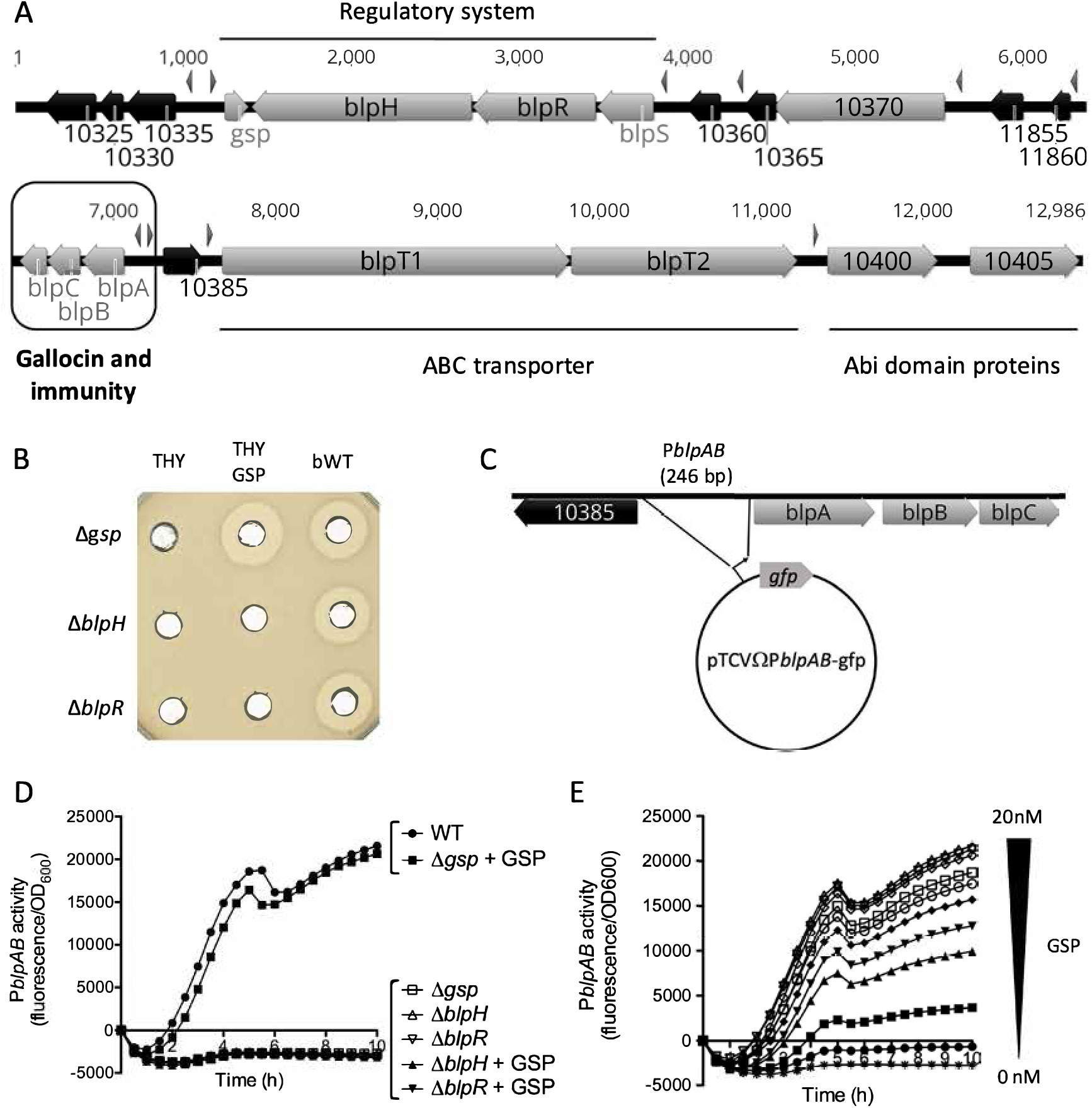
A three-component system activates gallocin transcription in *Sgg* UCN34. **A**: Gallocin locus in *Sgg* UCN34 (12,986 bp) extending from gene *gallo_rs10325* to *gallo_rs10405* (UCN34 genome ref seq: NC_013798.1, new annotation). Genes with a predicted function are indicated in grey, hypothetical genes are in black. Gene names are those given in this work or referred to the novel *“gallo_rs”* annotation (e.g.10325 for *gallo_rs10325*). Arrowhead above the genes indicates the presence of the conserved motif identified in **Fig. 4B**. **B**: Agar diffusion assay revealing *Sgg* UCN34 Δ*gsp*, Δ*blpH* and Δ*blpR* capacity to inhibit the growth of the gallocin-sensitive *Sgm* strain. Mutants were cultured either in THY or in THY supplemented with 20 nM of synthetic GSP (THY GSP). Activity of each bWT counterpart is also shown on the right. **C**: Schematic representation of the reporter plasmid pTCVΩP*blpAB-gfp* to monitor gallocin promoter activity. **D**: *PblpAB* activity (fluorescence divided by OD_600_) in *Sgg* UCN34 WT, Δ*gsp*, Δ*blpH* and Δ*blpR* in presence or absence of 20 nM of synthetic GSP. One representative curve of three independent experiments is shown here for each condition. **E**: *PblpAB* activity in *Sgg* UCN34 Δ*gsp* containing the reporter plasmid in presence of growing concentrations of synthetic GSP (curves from bottom to top were obtained in culture medium containing 0 to 20 nM GSP respectively, the concentration increasing by 2 nM between each curve. The three very similar upper curves were obtained with 16, 18 and 20 nM of GSP). One representative curve of three independent experiments is shown here for each condition.

To demonstrate that this regulatory system activates gallocin gene transcription, we constructed a reporter plasmid expressing *gfp* under the control of the gallocin operon promoter (pTCVΩP*blpAB-gfp*) to monitor the promoter activity by recording GFP fluorescence during growth (**Fig. 1C**). As shown in **Fig. S2**, *PblpAB* activity in *Sgg* UCN34 WT was null at the beginning of the culture, increased throughout growth and was maximal at the end of the exponential phase. Next, we showed that the *PblpAB* was completely inactive in *Streptococcus agalactiae* NEM316, which does not contain the specific regulatory system *gsp-blpRH* (**Fig. S2**). This result demonstrates that gallocin promoter activity depends on a *Sgg*-specific regulatory system. Consistently, the P*blpAB* promoter was found totally inactive in the three *Sgg* regulatory mutants Δ*gsp*, Δ*blpll*. and Δ*blpR* (**Fig. 1D**). Addition of increasing concentrations of GSP (2-20 nM) to the culture medium restored gallocin promoter activity in a dose-dependent manner in the *Sgg* Δ*gsp* mutant, but not in the Δ*blpH* and Δ*blpR* strains (**Fig. 1D and 1E**), confirming that GSP activates gallocin gene transcription through the BlpRH TCS. As shown in **Fig. 1E**, GSP is active at very low concentrations, since 16 nM of GSP was sufficient to fully activate gallocin promoter expression.

### 2 Identification of the regulon controlled by the 3-component regulatory system GSP / BlpHR in *S. gallolyticus*

In order to identify genes potentially involved in the production, maturation and secretion of gallocin, and to uncover new genes potentially co-regulated with gallocin genes, we performed a whole transcriptome analysis of the *Sgg* UCN34 WT, Δ*gsp*, Δ*blpH* and Δ*blpR* mutants. Total RNAs were extracted from exponentially growing *Sgg* cultures (THY OD_600_ 0.5). Overall comparison of the transcriptional profiles of these three mutants relatively to the parental UCN34 WT clearly shows that the main target of this regulatory system is the gallocin locus whose expression is strongly lowered in all three mutants (**Fig. 2A**). Selecting those genes whose transcription is significantly different (Log2 fold-change <-2 or >2; p-value<0.01) in at least one mutant as compared to *Sgg* WT UCN34 showed that 24 genes were downregulated in the three mutants (**Fig. 2B**).

**Figure 2:**
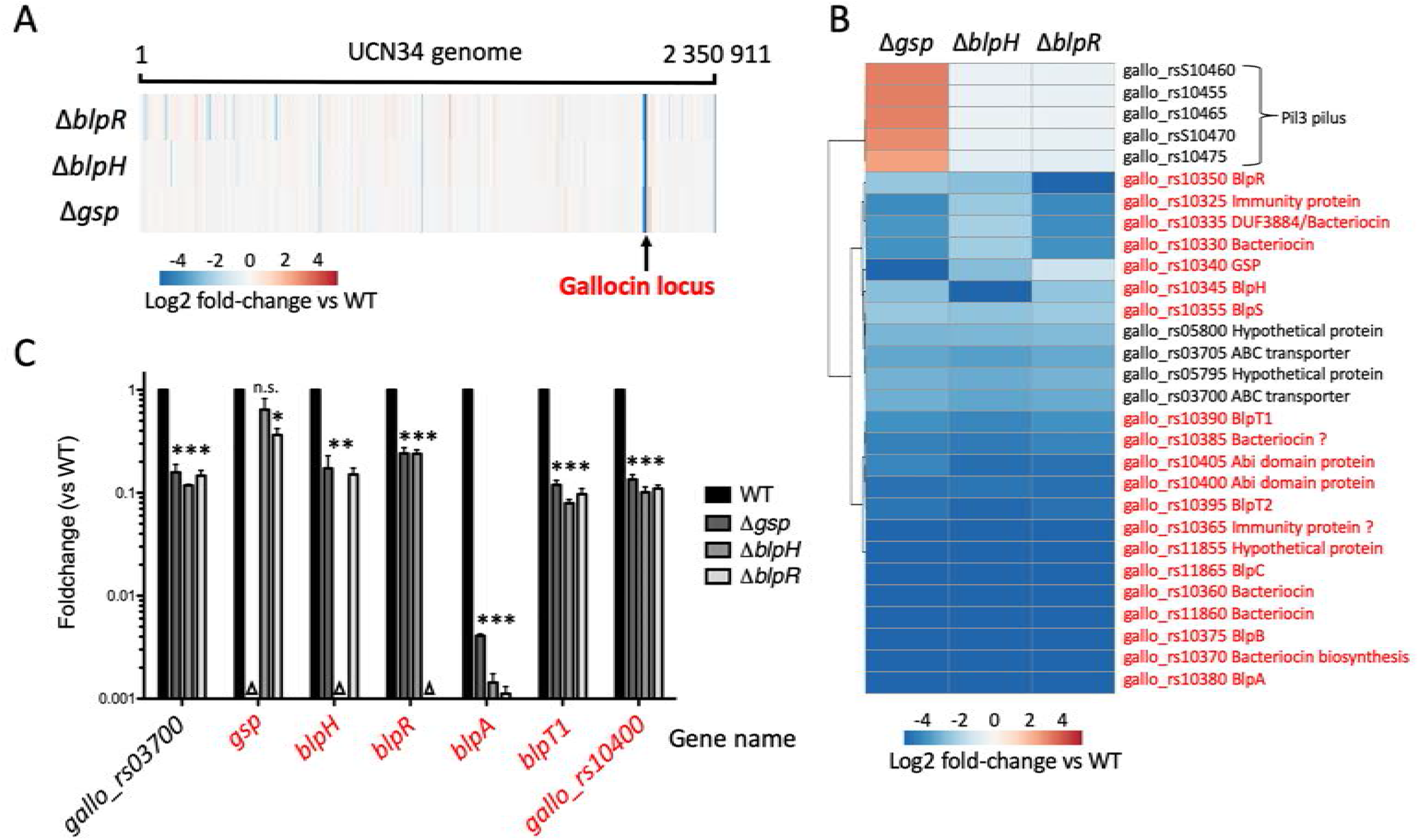
The whole regulon controlled by GSP/BlpHR. **A**: Heatmap representing the log2-foldchange in mRNA abundance (determined by whole transcriptomic analysis) of all the genes along the UCN34 genome in Δ*gsp*, Δ*blpH* and Δ*blpR* mutants compared to the parental *Sgg* UCN34 WT. **B**: Similar analysis as in A for selected genes whose log2-foldchange in mRNA abundance was significantly superior to 2 or inferior to −2 in at least one mutant (see details in material and methods). All the genes belonging to the gallocin locus are indicated in red with the corresponding gene product. Gene product was determined either with the genome annotation or by blast analysis. “?” indicates that the result of the blast analysis was not statistically significant. **C**: qRT-PCR data showing the foldchange in mRNA abundance in *Sgg* UCN34 Δ*gsp*, Δ*blpH* and Δ*blpR* compare to the WT. The identity of each mutant was confirmed by the absence of transcript indicated by Δ. Statistical analysis was performed to compare the expression of each gene in the three mutants compare to the WT by ANOVA (Significant difference was indicated with *: p-value<0,05; **: p-value<0,01; ***: p-value<0,001; n.s.: no significant difference)

Among these, 20 were part of the gallocin locus displayed in **Fig. 1A** (indicated in red in **Fig. 2B**): i) the regulatory module including *gsp*, *blpH*, *blpR*, plus the upstream adjacent *blpS* gene encoding a putative DNA binding protein; ii) the two genes encoding the gallocin peptides (*blpAB, gallo_rs10375/10380*) and the putative immunity protein (*blpC, gallo_rs11865y* iii) the two genes (*gallo_rs10390/10395*) encoding the ABC transporter whose role in the secretion of gallocin and GSP peptides is demonstrated in the accompanying paper (Harrington-Proutière et al., 2020); iv) *gallo_rs10370* which encodes a conserved protein with an undefined role in bacteriocin biosynthesis; v) *gallo_rs10400/10405* encoding Abi domain proteins; and vi) several hypothetical genes (indicated by black arrows in **Fig. 1A**). Sequence comparisons revealed that these hypothetical genes encode two putative bacteriocin operons (*gallo_rs10325/10335* and *gallo_rs10360/10365*) and two others single bacteriocins (*gallo_rs11860* and *gallo_rs10385*). Beside the gallocin locus, only four genes clustered in two different loci were found to be downregulated in the three regulatory mutants. These were the two adjacent genes *gallo_rs03700/03705* encoding a putative ABC transporter and *gallo_rs05795/05800* encoding hypothetical proteins of unknown function. Of note, the strong upregulation of the Pil3 pilus operon in the Δ*gsp* mutant (Fig. 2B) was shown to be due to a phase variation event as described previously (15).

To validate the transcriptome analysis, qRT-PCR was performed on 7 representative genes (**Fig. 2C**). qRT-PCR results confirmed the downregulation of these genes in absence of GSP or BlpRH TCS. Transcription of the core gallocin *blpABC* operon was more strongly reduced (>20-fold) as compared to the other genes of the locus such as those encoding the regulatory system (5-fold) and the ABC transporter (8-fold). It is worth noting that transcription of the *gsp* gene was only moderately altered in the Δ*blpH* and Δ*blpR* mutants as compared to the UCN34 WT (Fold-Change in Δ*blpH* : 0.159 by transcriptome and 0.64 by qRT-PCR; FC in Δ*blpR* : 0.4 by transcriptome and 0.37 by qRT-PCR). Together, these results show that this regulatory system strongly activates the transcription of several bacteriocin genes, including gallocin synthesis genes, and also induces its own transcription at a lower level.

### 3 Identification of a second regulator, BlpS, preventing transcriptional activation by BlpR

The *blpRH* genes encode a typical TCS composed of a response regulator BlpR that contain a CheY-homologous phospho-receiver domain and a LytTR DNA binding domain (**Fig. 3A**) and a sensor histidine kinase BlpH with 5 transmembrane regions. A second regulatory gene encoding a putative DNA-binding protein consisting entirely of a LytTR DNA-binding domain was found immediately upstream of *blpRH* **(Fig. 3A)**. We thus decided to test the role of this additional gene, designated *blpS* and likely co-transcribed with *blpRH*, in gallocin production. A clean in-frame deletion of this gene was performed in *Sgg* UCN34 to avoid polar effects on the downstream *blpRH* genes. Interestingly, the Δ*blpS* strain produced about 4-fold more gallocin than the WT as determined by serial dilution of the supernatant necessary to kill *Sgm* prey strain, suggesting that *blpS* represses gallocin gene expression. We then complemented the Δ*blpS* mutant and overexpressed *blpS* in the WT strain using a plasmid with an inducible promoter, pTCVΩP*tetO-blpS*. As a prerequisite, we first demonstrated that *PtetO* is functional in *Sgg* using pTCVΩP*tetO-gfp* as a reporter plasmid (**Fig S2**). We then showed that induction of *blpS* transcription leads to a decrease in gallocin production in both the Δ*blpS* and WT strains (**Fig. 3B**).

**Figure 3:**
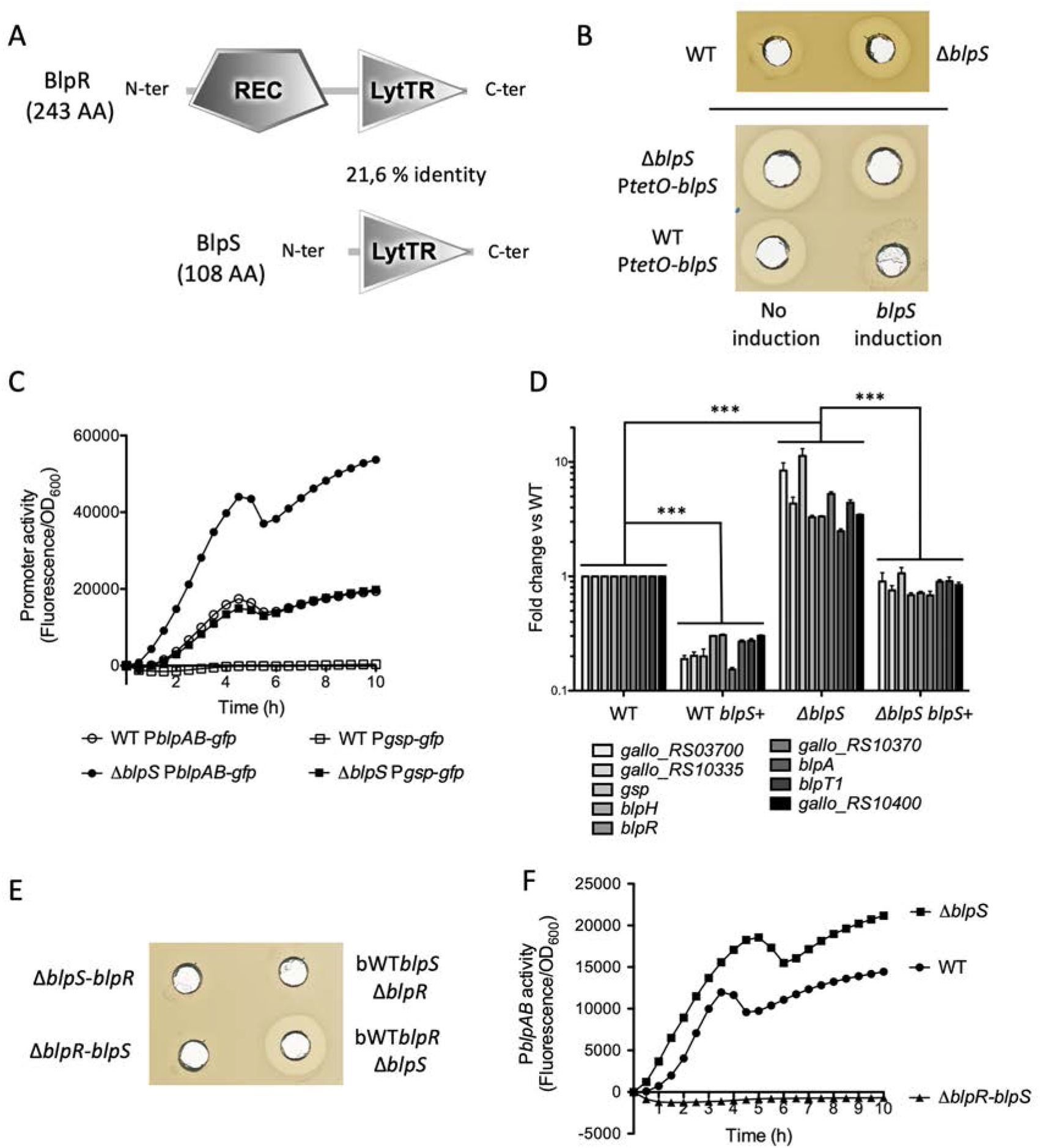
BlpS inhibits gallocin gene transcription. **A**: SMART domains identified in BlpR and BlpS proteins. REC stands for “cheY-homologous receiver domain” and LytTR for “LytTR DNA-binding domain”. The % of identity between the two LytTR domain was obtained using Geneious alignment. **B**: Agar diffusion assay showing gallocin activity in the culture supernatant against *Sgm*. Strain tested: *Sgg* UCN34 WT and Δ*blpS* (top part) and the same strains containing pTCVΩP*tetO*-*blpS* with or without induction of *blpS* expression with 200 ng/mL anhydrotetracycline (bottom part). **C**: Monitoring promoter activity of *PblpAB* (circle) and *Pgsp* (square) during growth in *Sgg* WT and Δ*blpS* mutant. One representative curve of three independent experiments is shown here for each condition. **D**: qRT-PCR data showing the foldchange in mRNA abundance between *Sgg* UCN34 pTCVΩP*tetO-blpS* (WT) and Δ*blpS pTCVΩPtetO-blpS (ΔblpS). blpS+* indicates the induction of *blpS* transcription with 200 ng/mL anhydrotetracycline. Statistical analysis was performed to compare the expression of each gene in the different conditions using ANOVA (*** indicates that results are significantly different with a p-value <0,0001) **E**: Agar diffusion assay to test gallocin activity in the culture supernatant of *Sgg* UCN34 Δ*blpS-blpR* (deletion of *blpS* in Δ*blpR*), Δ*blpR-blpS* (deletion of *blpR* in Δ*blpS*) and their respective bWT against *Sgm*. **F**: Monitoring *PblpAB* activity during growth in *Sgg* UCN34 WT, Δ*blpS* and Δ*blpR-blpS*. One representative curve of three independent experiments is shown here for each condition.

We next tested the effect of *blpS* deletion on the *blpAB* and *gsp* promoters. Reporter plasmids in which *gfp* expression was placed under the control of the *PblpAB* or *Pgsp* promoters were introduced in the *Sgg* WT and Δ*blpS* strains. As shown in **Fig. 3C**, expression from both *blpAB-* and *gsp-* promoters is strongly increased in the Δ*blpS* mutant as compared to the WT.

To determine the impact of BlpS on the whole regulon controlled by the GSP/BlpHR module, we determined by qRT-PCR the transcription levels of 9 different genes of this regulon, one located outside (*gallo_rs03700*) and eight within the gallocin genomic locus (*gallo_rs10335*, a putative bacteriocin, *gsp*, *blpH, blpR, gallo_rs10370, blpA, blpT1* and *gallo_rs10400*) in the WT and Δ*blpS* strains expressing *blpS* under the inducible promoter P*tetO*. Expression of the 9 tested genes was increased in the Δ*blpS* mutant compared to the WT strain in the absence of inducer (**Fig. 3D**). Induction of *blpS* expression reduced the transcript levels of the 9 tested genes in both the WT and Δ*blpS* strains (**Fig. 3D**). The highest fold-change between the WT and Δ*blpS* was observed for the *gsp* gene, which is upregulated more than 10-fold in the Δ*blpS* mutant. These results indicate that BlpS provides a negative feedback loop to control gallocin gene expression (**Fig. 3D**).

Finally, to determine the epistatic relationship of BlpR and BlpS on gallocin production, both genes were deleted, i.e. either by deleting the *blpS* gene in the Δ*blpR* mutant or vice versa. As shown in **Fig. 3E and F**, Δ*blpS-blpR* mutants were unable to produce gallocin and, consistently, the gallocin promoter P*blpAB* was totally inactive in these mutants. Together, these results demonstrate that BlpR is epistatic over BlpS.

### 4 Identification of a consensus DNA motif upstream from genes controlled by the BlpRH TCS

We next searched for conserved DNA motif(s) acting as putative binding site(s) in the promoter regions of the genes regulated by BlpR and BlpS. Our initial promoter sequence alignment of the gallocin locus genes (i.e. 250 base pairs upstream from the initiation codons) using Geneious software identified a conserved 15-bp motif (**Fig. 4A**). To improve the robustness of the 15-bp consensus sequence, promoters of the twelve putative operons regulated by BlpH/R were analyzed with the MEME software (http://meme-suite.org/tools/meme). A larger consensus sequence of 30-bp including most nucleotides of the previously identified 15-bp motif (12 bp out of 15) was identified in all regulated promoters (**Fig. 4B**). Mapping of this motif on the whole *Sgg* UCN34 chromosome showed that it is highly specific as it is present only upstream from the operons in the gallocin locus, as well as two other bicistronic loci, *gallo_rs03700* and *gallo_rs05800* identified by the transcriptome analysis (**Figs. 4C and 1A**). This 30-bp consensus motif, which is a likely binding site for BlpR and/or BlpS, contains 3 short repeats of 4 bp (C/TGAC). To map properly this motif in the P*blpAB* promoter, we determined the transcription start sites (TSS) of the gallocin *blpABC* operon by RACE-PCR. The 30-bp consensus motif lies just upstream of the −35 region of the operon promoter (**Fig. 4D**).

**Figure 4:**
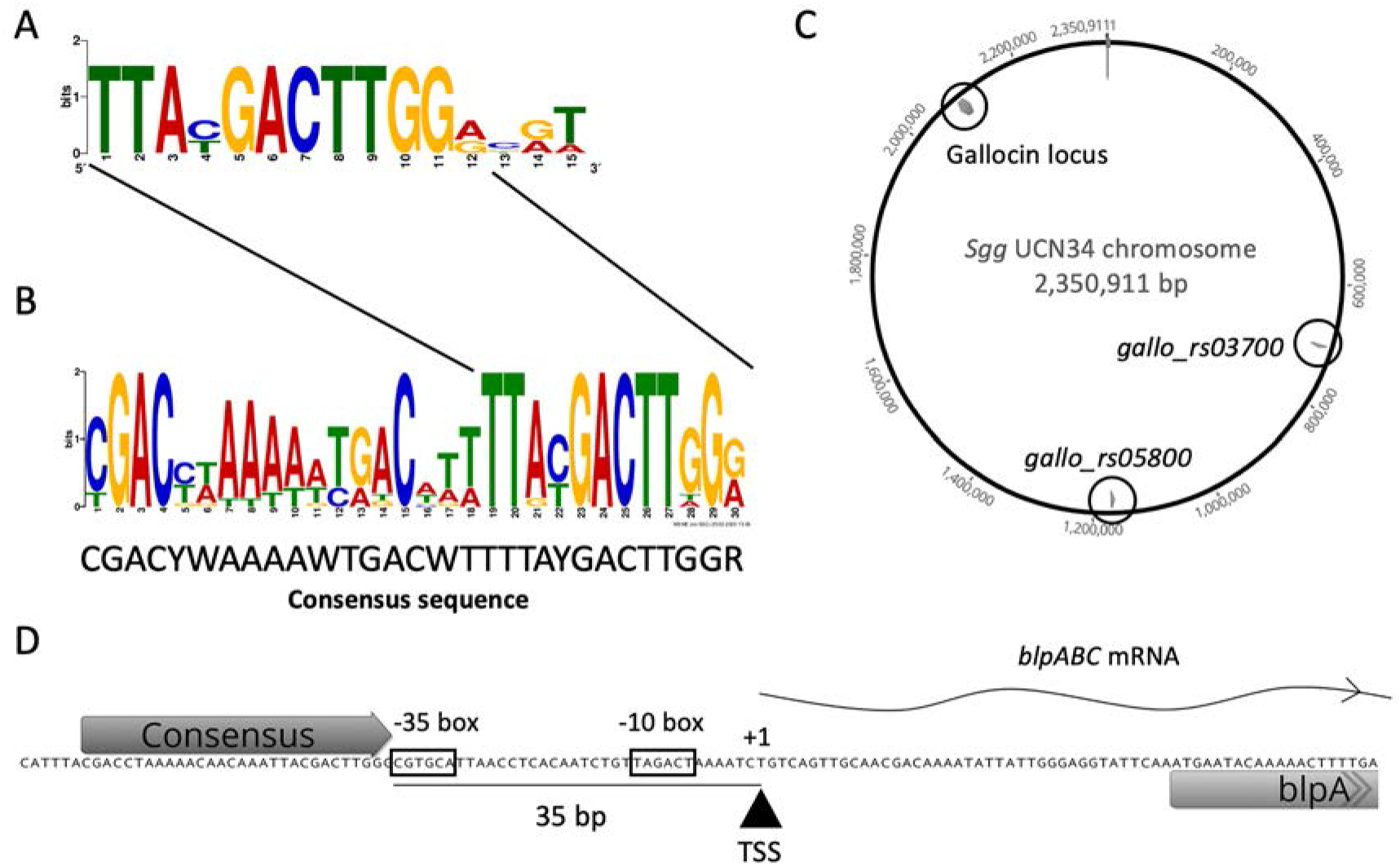
A conserved DNA motif is present upstream all the genes regulated by GSP-BlpHR. **A**: 15 bp DNA motif obtained by alignment of the promoters of the regulatory system, the bacteriocin accessory protein, gallocin genes, the ABC transporter and abi domain protein on https://weblogo.berkeley.edu/logo.cgi. **B**: A 30 bp consensus sequence identified by MEME in the twelve putative promoters regulated by BlpHR. The initial 15 bp motif is located at the 3’ end of this consensus motif. **C**: Mapping of the 30 bp consensus sequence on *Sgg* chromosome (with a maximum of 6 mismatches). The consensus sequences are represented by arrowheads and the name of the gene downstream the consensus is indicated. **D**: Determination of the transcription start site of *blpAB* and localization of the conserved motif. Putative −10 (TAGACT) and −35 (CGTGCA) promoter boxes were assigned based on the location of the (+1) transcription start site and the sequence of the canonical procaryotic promoter (TTGACA-X17-TATAAT).

### 5 Direct binding of BlpR and BlpS to various regulated promoters

To test the binding of BlpR and BlpS to the promoter regions that they control, electrophoretic mobility shift assays (EMSAs) were conducted on three regulated promoters *PblpAB, Pgsp*, and P*blpT*, and the unrelated P*gyrA* promoter as a negative control. BlpR and BlpS were produced as recombinant N-terminally histidine-tagged proteins in *Escherichia coli* BL21(DE3) and purified by immobilized metal-ion affinity chromatography. The purified proteins migrated around their expected molecular mass: 6xHis-BlpS: 14.1 kDa and 6xHis-BlpR: 29.3 kDa and were detected by Western blot using a His-tag monoclonal antibody (**Fig. S3A and B**). Of note, a second band of 25-30 kDa was identified for BlpS suggesting that this protein is able to form stable homodimers. Direct binding of recombinant BlpR and BlpS to the three regulated promoters, i.e. *PblpAB, PblpT1* and *Pgsp* was observed by EMSA in a dose-dependent manner while no binding to the control promoter P*gyrA* was detected (**Fig 5**).

**Figure 5:**
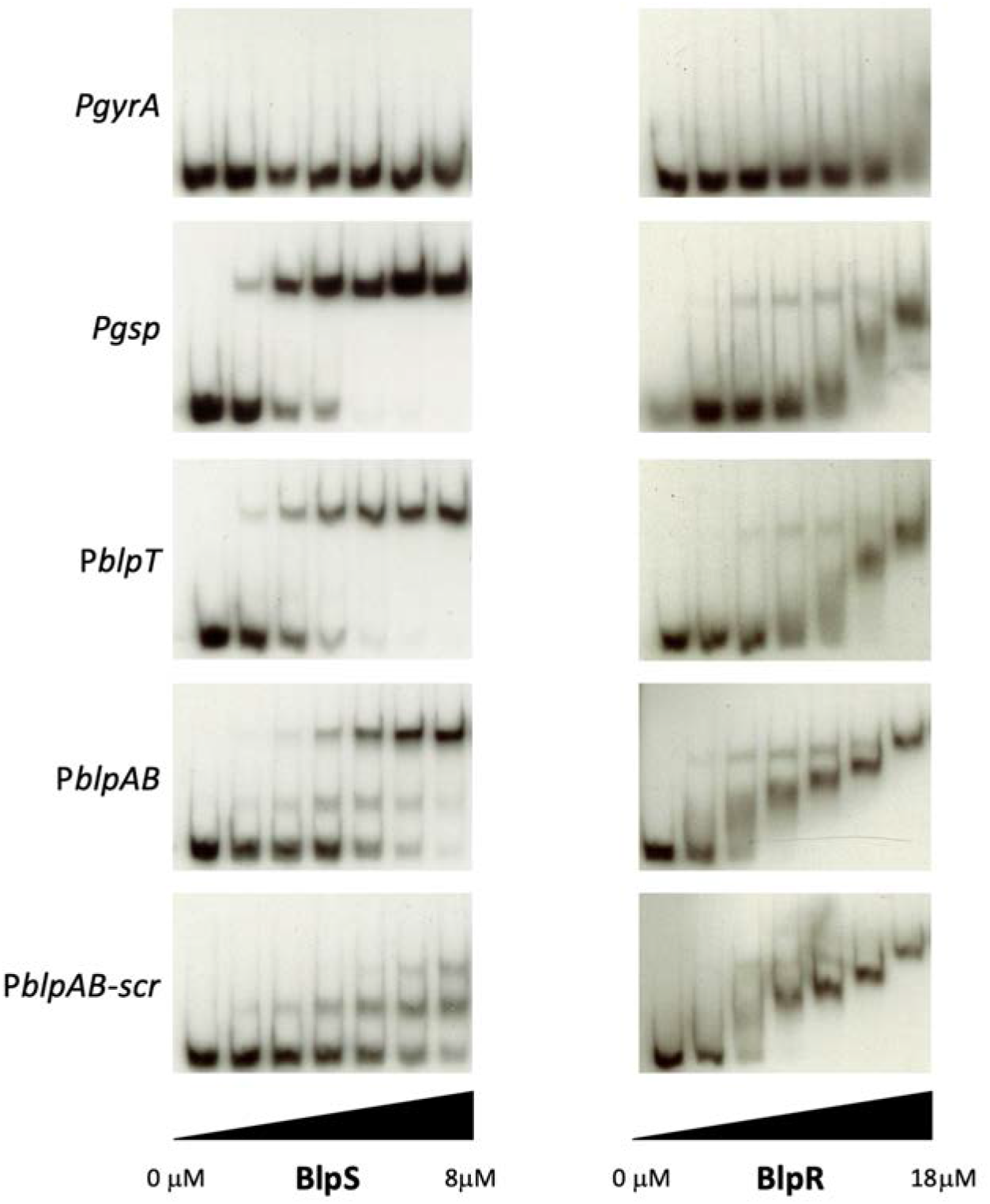
Binding of BlpR and BlpS to three regulated promoters. EMSA experiment demonstrating the binding of BlpS and BlpR to the promoter region of *gsp* (P*gsp*), *blpAB* (P*blpAB*), and *blpT* encoding the gallocin transporter. P*gyrA* was used as a negative control. Serial two-fold dilution of the protein (from right to left) were incubated with purified radiolabelled promoters before migration. The leftmost band correspond to migration of the promoter alone. The full sequences of the various promoters are indicated in Table 3. All these experiments were carried out in presence of 0,1μg/μL poly dI:dC to prevent aspecific binding of proteins to DNA. One representative EMSA of three independent experiments is shown here for each condition.

Finally, a similar EMSA experiment was performed on *PblpAB* where the initially identified 15-bp conserved motif was randomly scrambled (P*blpAB*-scr, **Fig. S4**). As shown in **Fig. 5**, binding of BlpS to the scrambled promoter was significantly altered while that of BlpR was unchanged.

## Discussion

*Streptococcus gallolyticus* subsp. *gallolyticus* (*Sgg*) belongs to Group D streptococci, a large group of phenotypically diverse bacteria known as the *S. bovis /S. equinus* complex (SBSEC), which consist of safe-graded bacteria used in food-fermentation, commensal bacteria of the gut and opportunistic pathogens in both humans and animals (16). *Sgg* is a commensal inhabitant of the rumen of herbivores, a complex ecological habitat harboring several thousand bacterial species. In humans, it is an opportunistic pathogen causing septicemia and endocarditis in elderly persons. Association between *Sgg* infections and underlying colon neoplasia has been reported by clinicians since the 1950s (2). Recently, we showed that *Sgg* strain UCN34 takes advantage of tumoral conditions to colonize the mouse colon (4). *Sgg* produces and secretes a specific bacteriocin, named gallocin, whose antimicrobial activity is potentiated by increased levels of secondary bile salts found in colonic neoplasia to inhibit the growth of closely related Enterococci commensals, thus creating a colonization niche for *Sgg* in tumoral conditions (4).

Gallocin is encoded by two genes, *blpA* and *blpB*, which are absent from closely related bacteria belonging to the SBSEC, including *Streptococcus gallolyticus* subsp. *macedonicus*. Gallocin is a class II bacteriocin, and members of this family are widespread among lactic acid bacteria, including streptococci. These molecules are usually directed against closely related bacteria competing within the same environment. The genetic locus encoding gallocin in *Sgg* UCN34 is complex and shares similarities with other prototypical class II bacteriocin loci with genes encoding a putative immunity protein, a dedicated ABC transporter, several other putative bacteriocins and a regulatory system (7, 17).

In this work, we demonstrated that gallocin production in *Sgg* is induced by a secreted peptide named GSP (for gallocin stimulating peptide) through the activation of a dedicated TCS composed of BlpH, a putative membrane histidine kinase, and BlpR, a putative cytoplasmic response regulator. Using a GFP-based reporter plasmid to monitor gallocin promoter (P*blpAB*) activity, we showed that synthetic 24-mer GSP activates gallocin promoter in a dose-dependent manner. GSP was shown to be secreted through the gallocin ABC transporter (designated BlpT). A structure/function analysis of the GSP peptide demonstrated the importance of its C-terminal half (Harrington-Proutière et al., accompanying paper). Since bacteriocin production has been linked to natural competence in various streptococci, including *S. pneumoniae*, *S. mutans, S. thermophilus*, we looked at competence induction in *Sgg* UCN34 using a reporter plasmid, in which the *comX* promoter was cloned upstream of the *gfp* and showed that *PcomX* was induced by XIP, the mature ComS peptide in agreement with previous result (18). However, no *PcomX* induction was observed using the GSP peptide (**Fig. S5**).

Our RNA-seq data revealed that transcription of five other putative bacteriocin genes was co-induced with gallocin genes, but whose roles are unknown. Only one of these has a double glycine motif in the N-terminal part, similar to the gallocin peptides while others have very different amino acid sequences (one being very rich in positively charged amino acids) suggesting that these other putative bacteriocins could have different roles. Only two additional operons encoding an ABC transporter and hypothetical proteins, located elsewhere in UCN34 genome, were co-induced with the gallocin locus.

We also uncovered the role of a second regulatory protein named BlpS which represses all the genes activated by GSP/BlpRH. This small protein of 108 amino acids consists almost entirely of a LytTR DNA-binding domain. Most proteins containing a LytTR domain studied previously also contain an additional phospho-acceptor domain typical of TCS regulators (Interpro domain : IPR007492, (19). Of note, two transcriptional regulators whose architecture is similar to that of BlpS were identified in *S. mutans*. However, these regulators are in operon with a transmembrane protein which inhibits their activity (20, 21). Thus, BlpS differs from these so-called LytTR Regulatory Systems (LRS) as it forms an operon with a classical TCS that it antagonizes. An *in silico* analysis revealed that 15,409 of the 80,096 LytTR-type regulators (Uniprot database) contained only this functional domain, and of these 1,565 have a size similar to BlpS (between 100 and 120 amino acids). These proteins were found both in gram-negative and gram-positive bacteria. Among them, we identified the homologous BlpS protein from *Streptococcus pneumoniae* belonging to the *blp* locus encoding pneumocins which is highly similar in its organization to the gallocin locus (22). We therefore propose that *blpS* gene of *S. pneumoniae* potentially encodes a negative regulator of pneumocin production.

To better understand the role of both BlpR and BlpS in regulation, we constructed the mutant Δ*blpR-blpS*. This mutant do not produce gallocin, showing that BlpR is necessary for transcriptional activation of gallocin genes even in the absence of the repressor BlpS. Then, we showed by EMSA that both BlpR and BlpS bound directly to the promoter region of the genes whose transcription is activated by BlpR. A conserved motif of 30-bp identified in these promoters could be the binding site of BlpR and BlpS. Thus, BlpS binding would antagonize the binding of BlpR to the regulated promoter to prevent its activation. EMSA experiments suggest that BlpS binds to the promoter sequences at lower concentrations than BlpR. As both regulators were produced in *E. coli*, the lower binding of BlpR could be due to the absence of phosphorylation by its cognate histidine kinase BlpH, while BlpS, lacking a regulatory domain, might be constitutively active. We postulate that, after phosphorylation by its dedicated histidine kinase BlpH, BlpR-P has a greater affinity for its DNA binding sites than its non-phosphorylated form and can thus replace BlpS.

A hypothetical model involving GSP-BlpRH-BlpS is proposed in **Fig. 6**. The gallocin promoter displays a poorly conserved −35 box which suggests that BlpR binding is necessary to recruit RNA-polymerase for ensuing transcription of gallocin genes. At low cell-density (i.e., no or very low amount of GSP), BlpS prevents the binding of the unphosphorylated BlpR to the promoters, thereby blocking transcription of the gallocin genes. At higher cell-density, sufficient amounts of GSP are present to induce BlpH-mediated phosphorylation of BlpR which, in turn, competes with BlpS to bind to the promoter region and trigger transcription of gallocin genes. Consistent with this model, deletion of BlpS results in expression of gallocin genes earlier in growth (Fig. 3C), suggesting that unphosphorylated BlpR is able to activate gallocin gene transcription in absence of BlpS.

**Figure 6:**
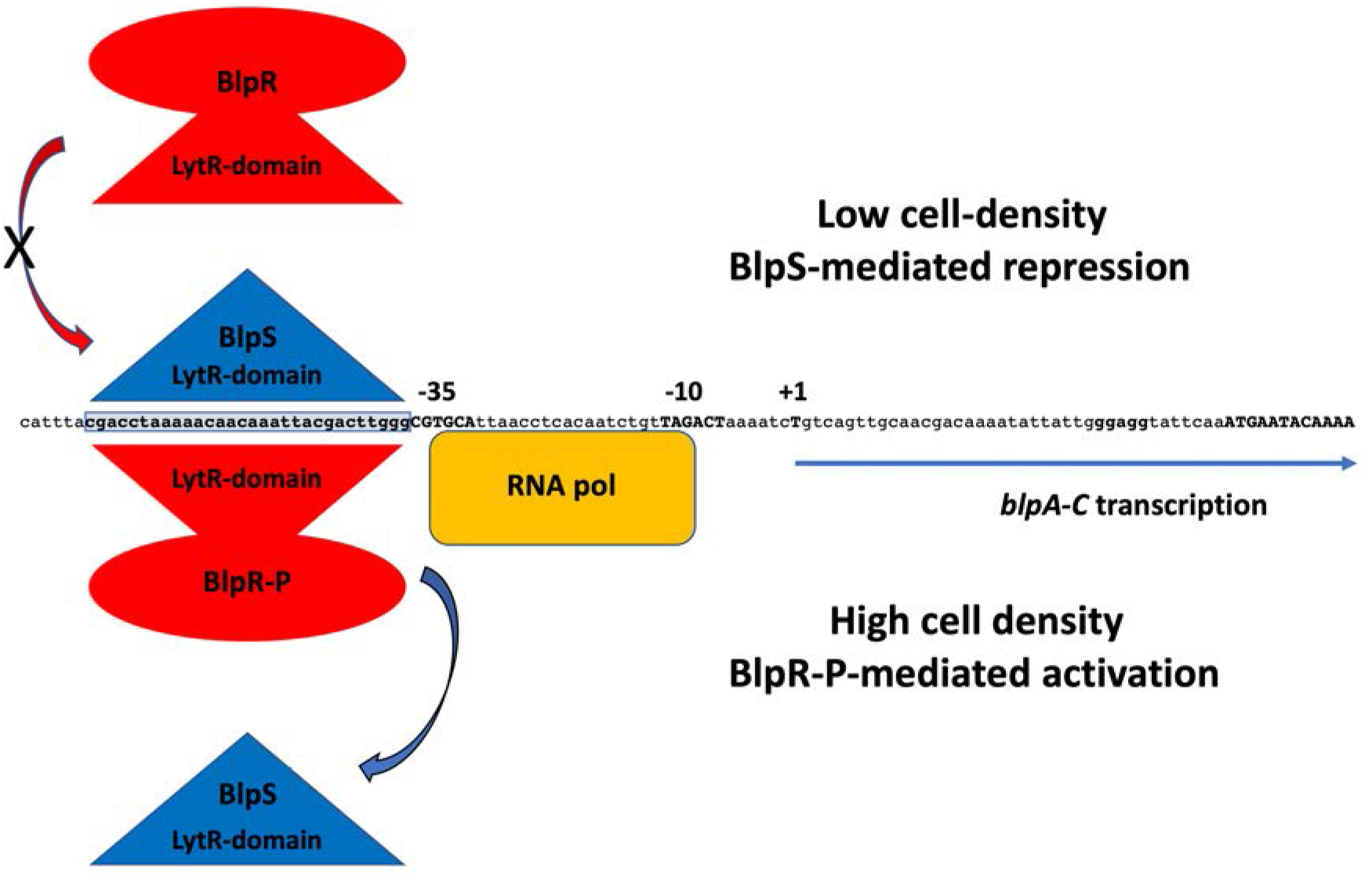
Hypothetical model of transcription regulation by BlpR and BlpS. At low cell-density, BlpS prevents BlpR binding to the conserved DNA motif found in the promoter of gallocin genes (*blpA-C*). At higher cell-density, GSP concentration is sufficient to induce BlpH-mediated phosphorylation of BlpR. Phosphorylated BlpR (BlpR-P) outcompetes BlpS and binds to the promoter of gallocin genes, resulting in RNA-polymerase recruitment and transcription of gallocin genes.

In conclusion, we identified here an atypical 4-component system involved in the regulation of bacteriocin production in *Sgg* UCN34, which could represent a new prototype of bacteriocin regulation. Bacteria have developed complex regulatory systems to control bacteriocin production in order to reduce its fitness cost. Indeed, we previously showed that the Δ*blp* mutant, which does not produce gallocin, colonize better than its Sgg WT counterpart in the non-tumoral murine intestinal tract. Although the Δ*blpS* mutant, which overproduced gallocin, did not exhibit a significant growth defect *in vitro* (data not shown), it remains possible that the increased production of gallocin could have an impact on its fitness *in vivo*, where nutrients are limited. Our hypothetical model propose that BlpS might be particularly important to prevent expression of gallocin at low cell-density which could result in the production of insuficient quantity of gallocin, favoring the appearance of gallocin-resistant strains.

Finally since gallocin is particularly active in tumoral conditions, it will be important in future studies to see if some tumoral metabolites could induce gallocin transcription *in vivo*.

## Material and Methods

### Cultures, bacterial strains, plasmids and oligonucleotides

*Streptococci* used in this study were grown at 37°C in Todd-Hewitt broth supplemented with yeast extract 0.5% (THY) in standing filled flasks. When appropriate, 10μg/mL of erythromycin were added to the culture for plasmid maintenance.

Plasmid construction was performed by: PCR amplification of the fragment to insert in the plasmid with Q5^®^ High-Fidelity DNA Polymerase (New England Biolabs), digestion with the appropriate FastDigest restriction enzyme (ThermoFisher), ligation with T4 DNA ligase (New England Biolabs) and transformation in commercially available TOP10 competent *E. coli* (ThermoFisher). *E. coli* transformants were cultured in LB Miller’s supplemented with 150 μg/mL erythromycin (for pG1derived plasmids) or 50 μg/mL kanamycin (pTCV-derived and pET28a plasmids). Verified plasmids were electroporated in *S. agalactiae* NEM316 and mobilized from NEM316 to *Sgg* UCN34 by conjugation as described previously (23). All the strains used and constructed in this study are listed in table 1, primers in table 2.

**Table 1:**
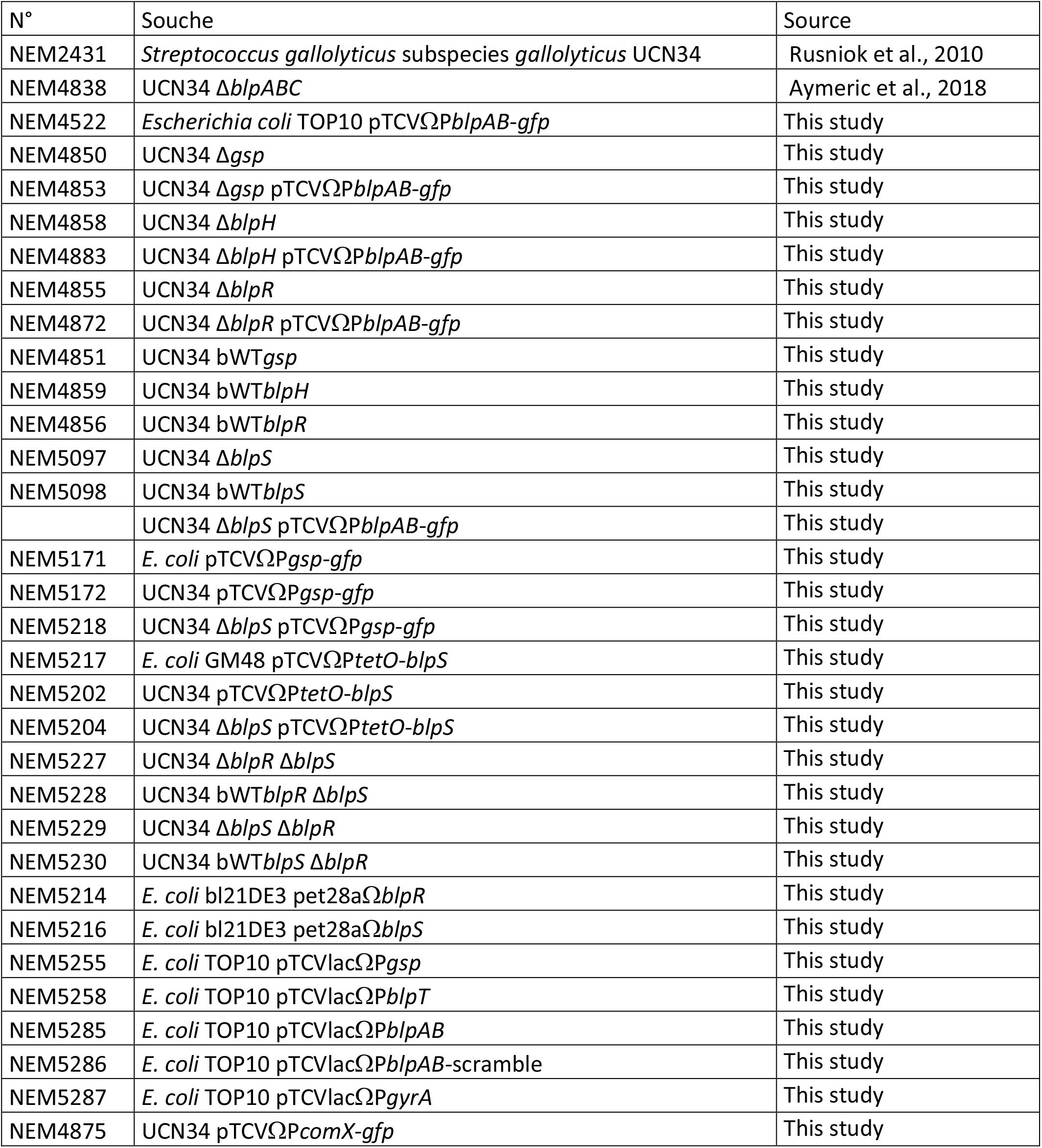
Strains used in this study

| N° | Souche | Source |
| --- | --- | --- |
| NEM2431 | *Streptococcus gallolyticus* subspecies *gallolyticus* UCN34 | Rusniok et al., 2010 |
| NEM4838 | UCN34 $\Delta blpABC$ | Aymeric et al., 2018 |
| NEM4522 | *Escherichia coli* TOP10 pTCVΩPblpAB-gfp | This study |
| NEM4850 | UCN34 $\Delta gsp$ | This study |
| NEM4853 | UCN34 $\Delta gsp$ pTCVΩPblpAB-gfp | This study |
| NEM4858 | UCN34 $\Delta blpH$ | This study |
| NEM4883 | UCN34 $\Delta blpH$ pTCVΩPblpAB-gfp | This study |
| NEM4855 | UCN34 $\Delta blpR$ | This study |
| NEM4872 | UCN34 $\Delta blpR$ pTCVΩPblpAB-gfp | This study |
| NEM4851 | UCN34 bWTgsp | This study |
| NEM4859 | UCN34 bWTblpH | This study |
| NEM4856 | UCN34 bWTblpR | This study |
| NEM5097 | UCN34 $\Delta blpS$ | This study |
| NEM5098 | UCN34 bWTblpS | This study |
| | UCN34 $\Delta blpS$ pTCVΩPblpAB-gfp | This study |
| NEM5171 | *E. coli* pTCVΩPgsp-gfp | This study |
| NEM5172 | UCN34 pTCVΩPgsp-gfp | This study |
| NEM5218 | UCN34 $\Delta blpS$ pTCVΩPgsp-gfp | This study |
| NEM5217 | *E. coli* GM48 pTCVΩPtetO-blpS | This study |
| NEM5202 | UCN34 pTCVΩPtetO-blpS | This study |
| NEM5204 | UCN34 $\Delta blpS$ pTCVΩPtetO-blpS | This study |
| NEM5227 | UCN34 $\Delta blpR \Delta blpS$ | This study |
| NEM5228 | UCN34 bWTblpR $\Delta blpS$ | This study |
| NEM5229 | UCN34 $\Delta blpS \Delta blpR$ | This study |
| NEM5230 | UCN34 bWTblpS $\Delta blpR$ | This study |
| NEM5214 | *E. coli* bl21DE3 pet28aΩblpR | This study |
| NEM5216 | *E. coli* bl21DE3 pet28aΩblpS | This study |
| NEM5255 | *E. coli* TOP10 pTCVlacΩPgsp | This study |
| NEM5258 | *E. coli* TOP10 pTCVlacΩPblpT | This study |
| NEM5285 | *E. coli* TOP10 pTCVlacΩPblpAB | This study |
| NEM5286 | *E. coli* TOP10 pTCVlacΩPblpAB-scramble | This study |
| NEM5287 | *E. coli* TOP10 pTCVlacΩPgyrA | This study |
| NEM4875 | UCN34 pTCVΩPcomX-gfp | This study |

**Table 2:**
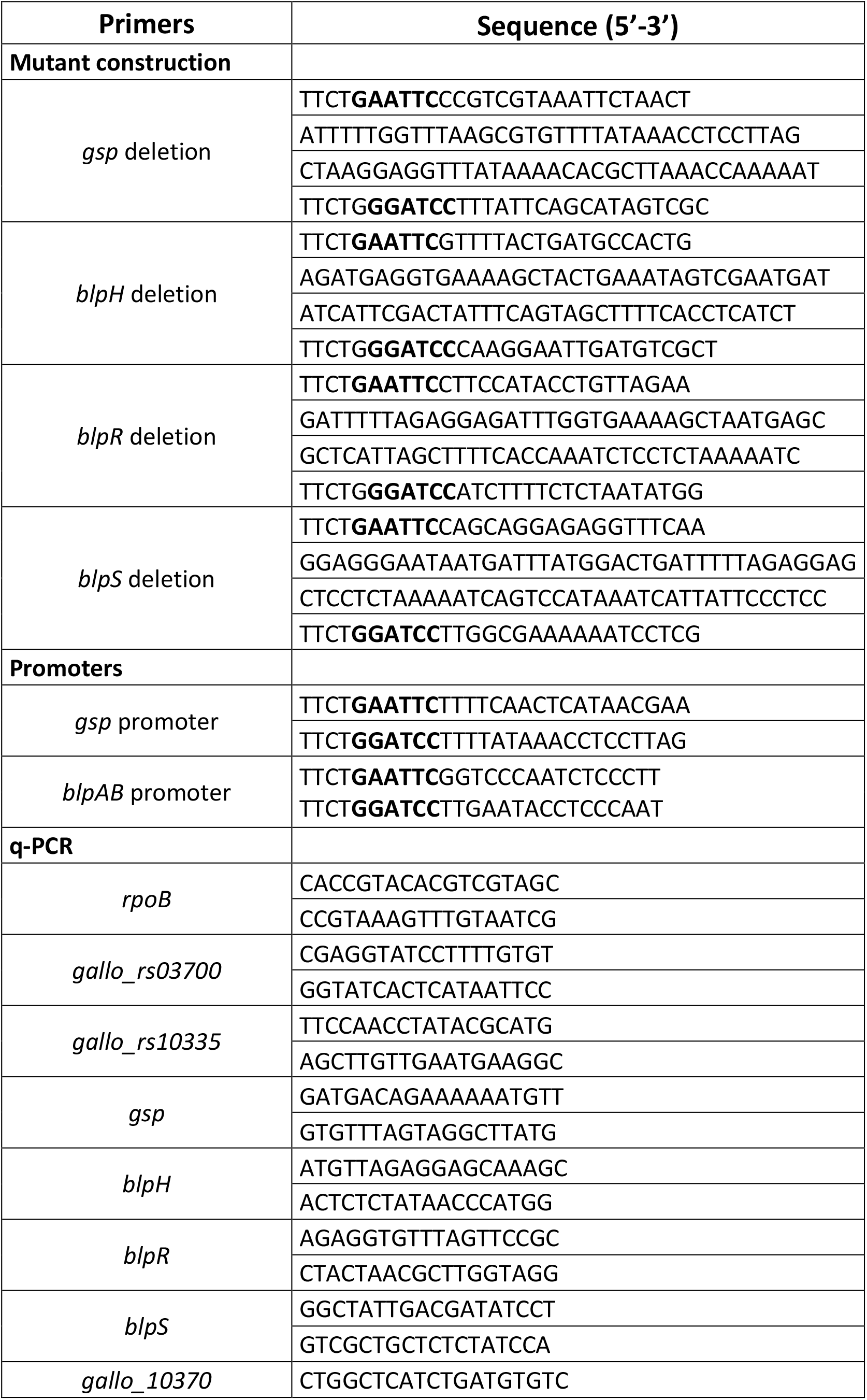
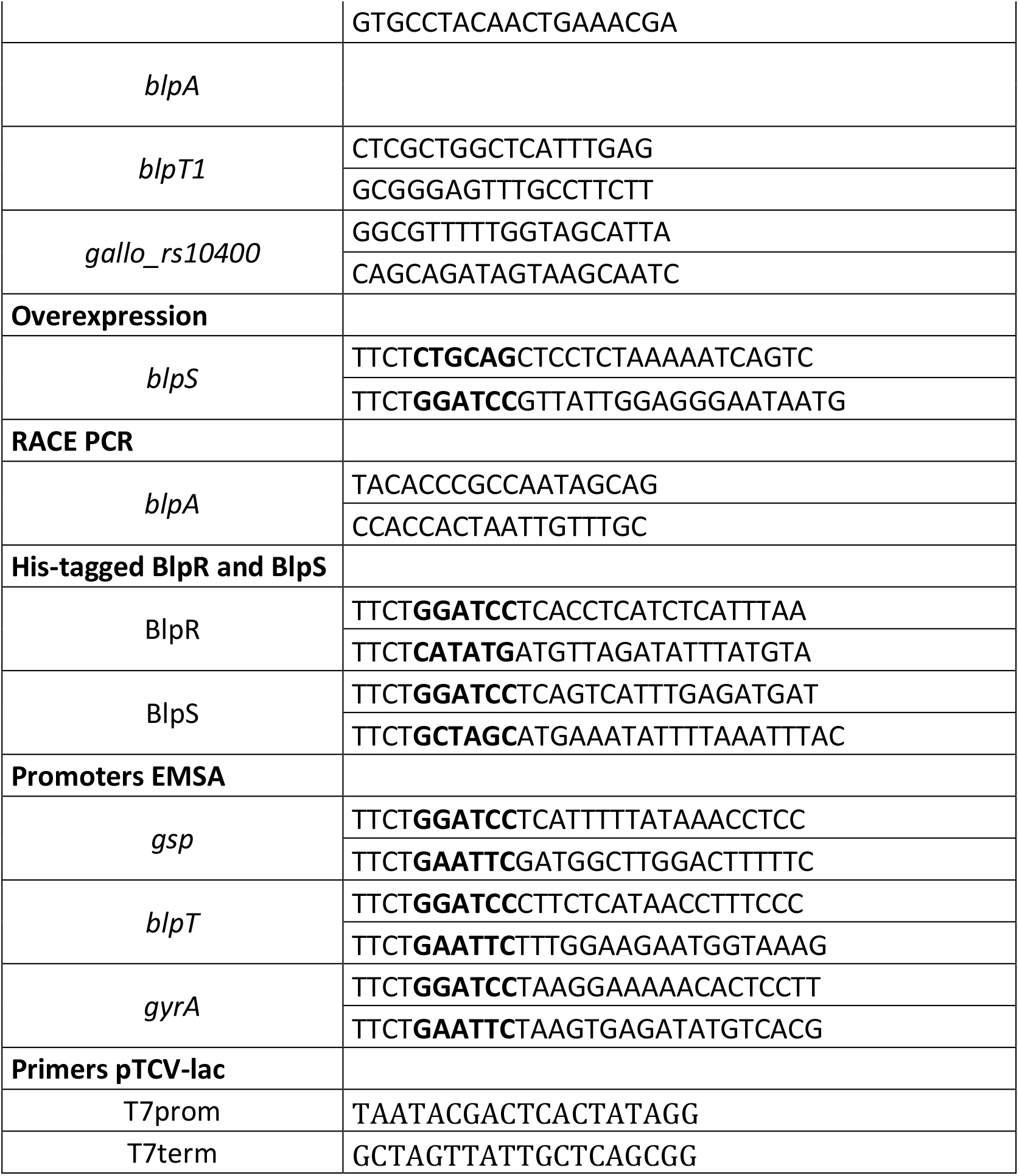
Primers used in this study. Restriction sites are indicated in bold

### Construction of markerless deletion mutants in Sgg UCN34

In frame deletion mutants were constructed as described previously (23). Briefly, the 5’ and 3’ region flanking the region to delete were amplified and assembled by splicing by overlap extension PCR and cloned into the thermosensitive shuttle vector pG1. Once transformed in UCN34, the cells were cultured at 38°C with erythromycin to select for the chromosomal integration of the plasmid by homologous recombination. About 4 single cross-over integrants were serially passaged at 30 °C without antibiotic to facilitate the excision of the plasmid resulting either in gene deletion or back to the WT (bWT). In-frame deletions were identified by PCR and confirmed by DNA sequencing of the chromosomal DNA flanking the deletion.

### Gallocin production assays

Briefly, one colony of the indicator strain, here *Streptococcus macedonicus*, was resuspended in 2 mL THY, grown until exponential phase, poured on a THY agar plate and the excess liquid was removed and left to dry under the hood for about 20 min. Using sterile tips, 5-mm-diameter wells were dug into the agar. Each well was then filled with 80 μL of filtered supernatant from 5 h cultures (stationary phase) of *Sgg* WT or mutant strains and supplemented with Tween 20 0.1% final concentration. Inhibition rings around the wells were observed the following morning after overnight incubation at 37°C.

### Monitoring promoter activity using a fluorescent reporter

Promoter sequences of genes encoding gallocin (*PblpAB*) or GSP (P*gsp*) were amplified with overhanging *Eco*R1 and *BamH1* sites and cloned in the reporter pTCVΩgfp vector upstream from the *gfp* gene to control its expression. Bacteria containing the plasmid were inoculated at an initial OD_600_ of 0.1 from fresh agar plates in 200 μL of media in 96-well black plates. Due to the high autofluorescence of the THY medium, we switched to M9 medium supplemented with yeast extract 0.5% and glucose 0.2% (M9Y). Promoter activity was then followed by continuous measurement of the growth and GFP fluorescence (one measure every 30 min during 10 h) of the cultures with the Synergy™2 Multi-detection microplate reader (Biotek). Promoter activity was then estimated by dividing the fluorescence value by the OD_600_ value for each time point.

### Induction of *PtetO* promoter in *Sgg*

Using the reporter plasmid pTCVΩP*tetO-gfp*, we defined the minimal concentration of anhydrotetracycline necessary to fully induce *PtetO* promoter in *Sgg* UCN34 to be 200 ng/mL. The *blpS* gene was cloned in the pTCV-*PtetO* in *E. coli* and then introduced in WT *Sgg* UCN34 and Δ*blpS* and *blpS* expression was induced with 200 ng/mL anhydrotetracycline.

### Transcriptomic analysis and Real-Time Quantitative Reverse Transcription

Total RNAs were extracted from exponentially growing *Sgg* strains (OD_600_=0.5) in THY at 37°C with the MP Biomedicals™ FastRNA™ Pro Blue Kit following the manufacturer’s recommendations. 20 μg of bacterial RNA were treated with DNase I (Invitrogen™ Ambion™ TURBO DNA-free Kit) to remove residual genomic DNA and then DNase I was inactivated with the recommended reagent.

For whole transcriptomic analysis, rRNA were depleted from 0.5 μg of total RNA using the Ribo-Zero rRNA Removal Kit (Bacteria) from Illumina. Sequencing libraries were constructed using the TruSeq Stranded mRNA Sample preparation kit (20020595) following the manufacturer’s instructions (Illumina). The directional libraries were controlled on Bioanalyzer DNA1000 Chips (Agilent Technologies) and concentrations measured with the Qubit^®^ dsDNA HS Assay Kit (ThermoFisher). Sequences of 65 bases were generated on the Illumina Hiseq 2500 sequencer. Reads were cleaned of adapter sequences and low-quality sequences using cutadapt version 1.11 (24). Only sequences at least 25 nt in length were considered for further analysis. Bowtie version 1.2.2 (25), with default parameters, was used for alignment on the reference genome (NC_013798.1, from NCBI). Genes were counted using featureCounts version 1.4.6-p3 (26) from Subreads package (parameters: -t gene -g locus_tag - s 1). Count data were analyzed using R version 3.5.1 and the Bioconductor package DESeq2 version 1.20.0 (27). The normalization and dispersion estimation were performed with DESeq2 using the default parameters and statistical tests for differential expression were performed applying the independent filtering algorithm. A generalized linear model was set in order to test for the differential expression between the WT, Δ*gsp*, Δ*blpH* and Δ*blpR* biological conditions. For each pairwise comparison, raw p-values were adjusted for multiple testing according to the Benjamini and Hochberg (BH) procedure (28) and genes with an adjusted p-value lower than 0.05 were considered differentially expressed. The raw data of this transcriptomic analysis are available on GEO data server (accession number: GSE148401).

The heatmaps representing the results in this paper were realized with all the genes from *Sgg* UCN34 genomes (figure 2A) or with genes whose expression is significantly different in at least one mutant compare to the WT (figure 2B). Here, gene expression was considered significantly different if the adjusted p-value was lower than 0.01 and if the log2 fold-change in gene expression was inferior to −2 or superior to 2 compare to WT. Finally, some non-assigned genes whose expression was very low (50-150 reads per gene) but significantly different in the Δ*blpR* mutant were also suppressed from this heat-map for clarity purpose.

For real-time quantitative reverse transcription, cDNAs were obtained from 1 μg of RNA treated with DNase I using the iScript™ cDNA Synthesis Kit. Real-time quantitative PCR was carried out on three independent biological replicates in a CFX96 Touch™ Real-Time PCR Detection System (Bio-Rad) in 20 μL mix containing 10 μL EvaGreen Universal qPCR Supermix (Bio-Rad), 1 μL gene specific primers (10 μM) and 5 μL of a 100-fold dilution of cDNA. Fold-change in expression compare to WT were obtained by the 2^-ΔΔCt^ method. For statistical analysis, RT-qPCR data were analyzed using ANOVA: for each gene a model that explains ΔCt values was fitted, including the replicate effect as random. The model also includes the strain (Fig. 2C) or the strain, the condition and their interactions (Fig. 3D) as fixed effects. Pairwise comparisons were tested thanks to the emmeans R package version 1.4.2 and p-values were adjusted for multiple testing using the Tukey method.

### RACE-PCR

RACE-PCR to determine the transcriptional start site was performed with the 5’ RACE System for Rapid Amplification of cDNA Ends (ThermoFisher) following the manufacturer protocol. Briefly, total RNAs were purified from a *Sgg* UCN34 WT culture as indicated above. cDNA of *blpABC* mRNA was obtained by reverse transcription with a gene specific primer. A homopolymeric tail was added to the 3’-end of the cDNA, corresponding to the former 5’ end of the mRNA. The cDNA was amplified by PCR with another gene specific primer located in the cDNA and a deoxyinosine-containing anchor primer provided in the kit that anneals to the homopolymeric tails of the cDNA. The resulting PCR fragment was cloned in the Zero Blunt™ TOPO™ plasmid (Thermofisher) and transformed in *E. coli*. After purification, plasmids were sequenced and sequence alignment was performed to identify the transcription start site.

### Production and purification of His-tagged recombinant proteins

Full length *blpR* and *blpS* were cloned in the pET28a vector in order to obtain 6His-tagged proteins at their N-terminus and after sequence verification the recombinant plasmids were transferred in the host expression vector *E. coli* BL21(DE3). Histidine-tagged proteins were purified as previously described (29). Briefly, *E. coli* cells carrying the plasmid were grown in 500 mL LB supplemented with kanamycin (50 μg/mL) at 37°C with agitation until reaching OD_600_~ 0.5. At this point, 1mM IPTG was added to the culture to induce protein expression and the culture was incubated for 3 h at 37°C with agitation. Cells were pelleted, incubated overnight at −20°C and resuspended in 20 mL of basis buffer (29) containing 1 mg/mL of lysozyme to lyse them. Cell debris were eliminated by centrifugation and 1 mL of Ni-NTA superflow beads (Qiagen) was added to bind his-tagged proteins. After washing on a gravity flow column, his-tagged proteins were eluted with an elution buffer containing 500 mM imidazole (29). Fraction containing the recombinant protein were pooled and resuspended in the following buffer (NaH_2_PO_4_ 50mM, NaCl 300mM, DTT 1mM, glycerol 20%, pH=8) using PD10 columns. Purified proteins were conserved at −80°C. Just before utilization, proteins were concentrated around 10-fold on Vivaspin column (5kDa cutoff) and protein concentration was estimated by measurement of the OD_280_.

### DNA-protein interactions

Electrophoretic mobility shift assay (EMSA) was performed as described previously (30). Briefly, promoter sequences (Table 3) of about 150 bp were amplified by PCR and cloned in the pTCV-lac vector (31). *blpAB* promoter and its scrambled derivative were chemically synthesized by Genecust and cloned in pTCV-lac vector. All promoters were then amplified by PCR with radiolabeled primer specific for the plasmid cloning site (T7prom, T7term). Radiolabeled PCR diluted 100-fold were incubated 30 min in binding buffer (25 mM Na_2_HPO_4_/NaH_2_PO_4_ pH 8, 50 mM NaCl, 2 mM MgCl_2_, 1 mM DTT, 10% glycerol) supplemented with 0.02 μg / μl BSA and 0.1 μg / μl of Poly DI-DC (Sigma) in presence of serial dilution of purified BlpR/BlpS or buffer. After migration of the different reactions on a 6% polyacrylamide gel for 1 h, gels were analyzed by autoradiography.

**Table 3:**
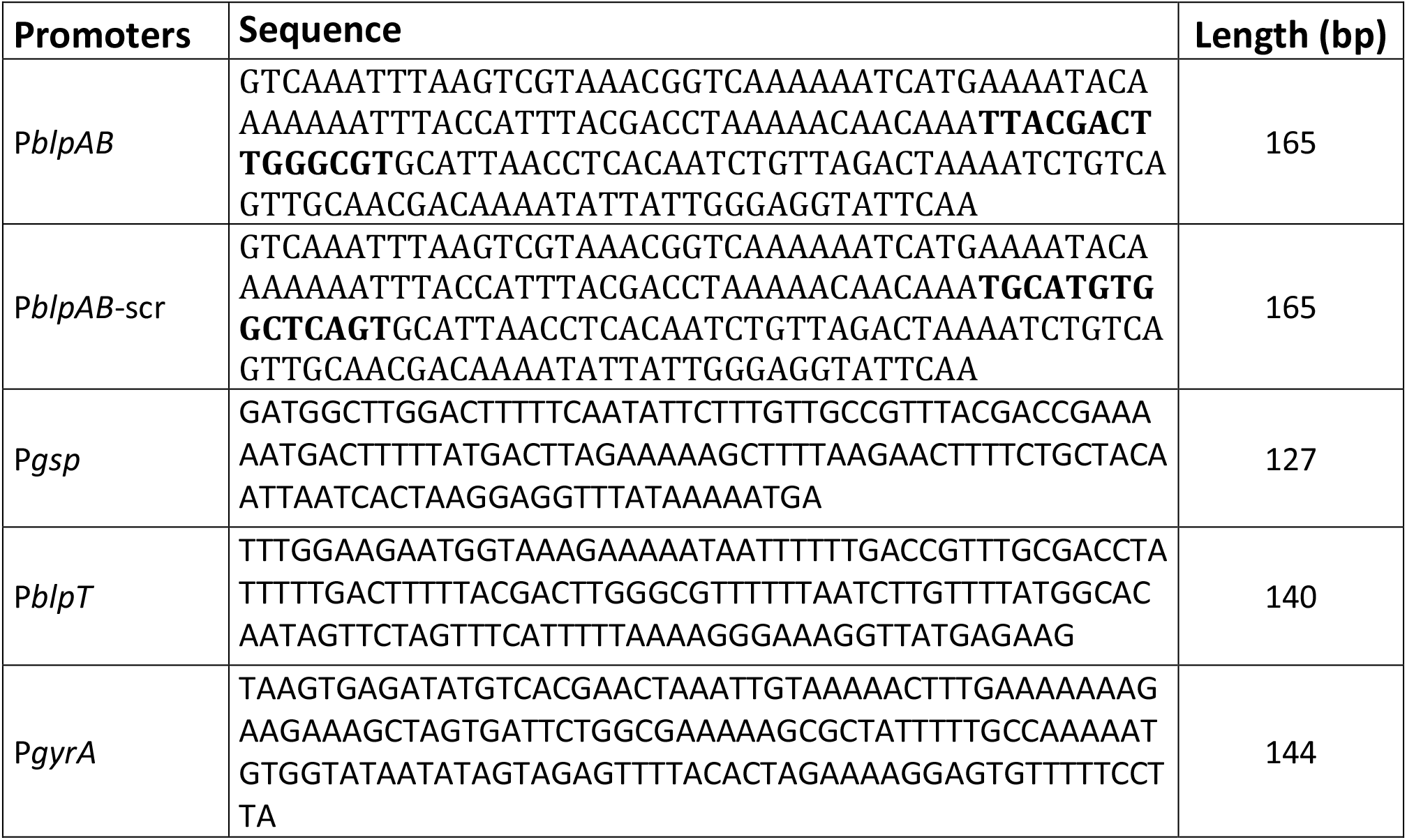
Sequence of the promoters tested by EMSA. The *PblpAB* sequence that is scrambled in *PblpAB-scr* is indicated in bold.

## Acknowledgements

We would like to thank particularly Odile Sismeiro for the construction of the transcriptome library and Rachel Legendre for the bioinformatic analysis of the transcriptome and its deposit on GEO database. We are also grateful to Christophe Rusniok for his help in the search of LytTR-only proteins, Christine Dang for the reporter plasmid construction and Tarek Msadek for careful and critical reading of the manuscript. We also thank Myriam Gominet and Maria Vittoria Mazzuoli for providing their expertise in EMSA experiments and Gaël Millot for its help with the statistical analysis of qRT-PCR data. This study has been funded by the Institut National contre le Cancer (INCA) PLBIO 16-025 attributed to S. Dramsi.

## Supplementary figures legend

**Sup Fig 1:**
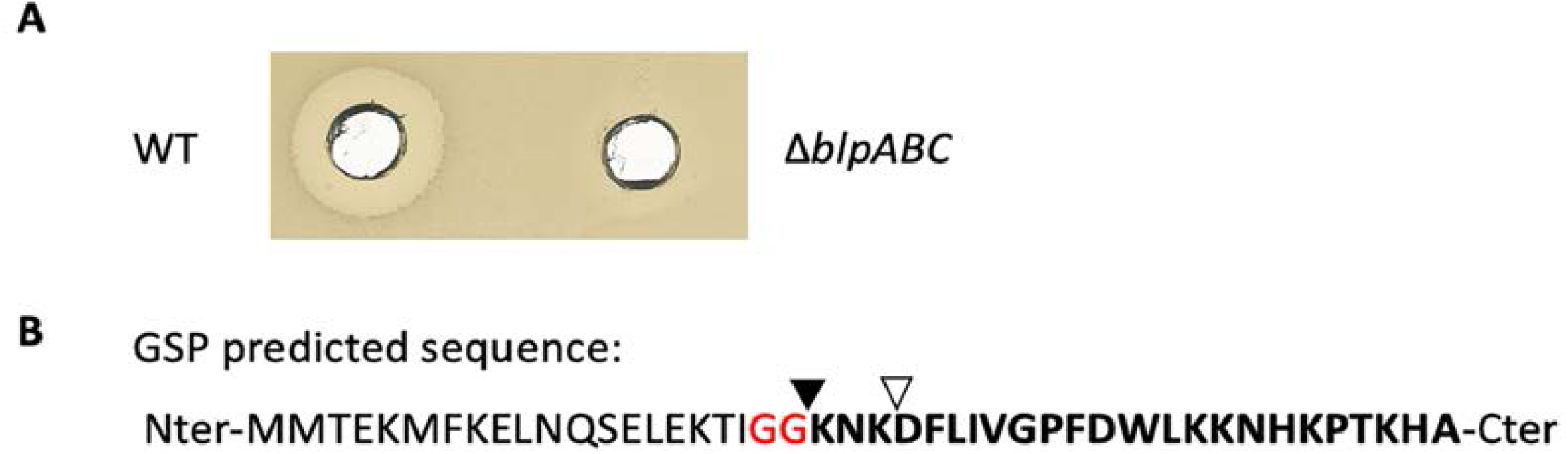
Susceptibility of *Streptococcus gallolyticus* subsp. *macedonicus* to gallocin and GSP amino acid sequence. **A**: Agar diffusion assay of *Sgg* UCN34 WT and Δ*blpABC* supernatant on a lawn of *Sgm*. **B**: Amino acid sequence of GSP obtained by translation of *gsp* gene. Predicted cleavage site indicated by a black arrowhead occurs after the double glycine motif in red. The synthetic peptide used in this study is composed of the 24 C-terminal residues in bold. The white arrowhead indicates the cleavage site identified after GSP purification from culture supernatant and sequencing (Harrington et Tal-Gan 2018; Harrington-Proutière et al., 2020, companion paper).

**Sup Fig 2:**
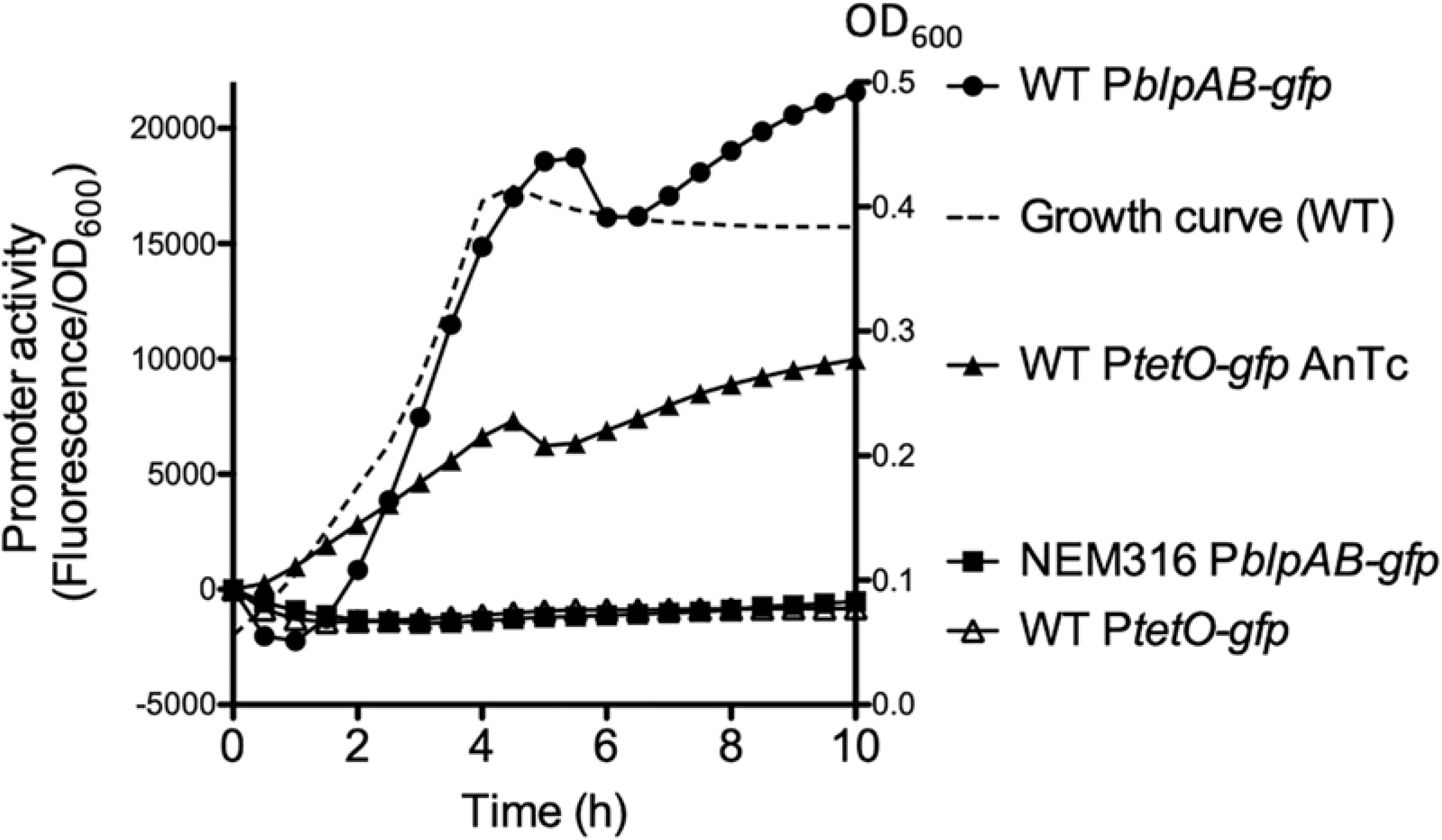
Monitoring GFP fluorescence using *PblpAB-* or PtetO-inducible promoters. *PblpAB* is active in *Sgg* UCN34 WT but not in GBS NEM316. P*tetO* was induced with 200 ng/mL of anhydrotetracycline (AnTc) in *Sgg* UCN34. All the strains exhibited very similar growth. For clarity purpose, only the growth curve of *Sgg* WT (pTCVΩ*PblpAB-gfp*) is presented here as a dotted curve. One representative curve of three independent experiments is shown here for each condition.

**Sup Fig 3:**
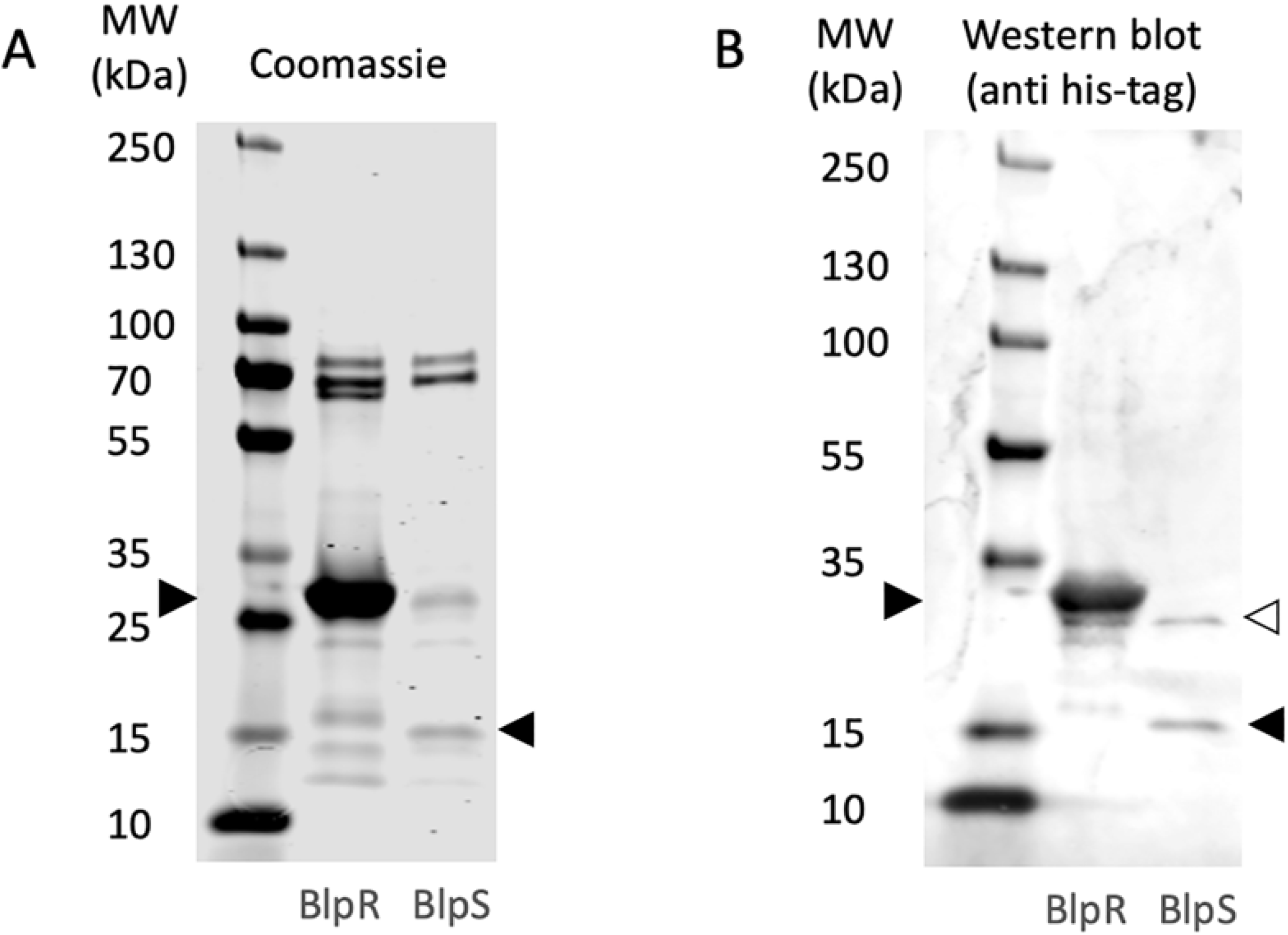
SDS-PAGE and Western-blot analysis of purified histidine-tagged recombinant BlpR and BlpS proteins. **A**: Visualization of purified recombinant BlpR and BlpS 10-fold concentrated on Vivaspin column (cut off 5,000 Da) by Coomassie staining and **B**-Western blotting using primary anti His tag monoclonal antibody followed by a secondary fluorescent conjugated goat anti mouse antibody. Position of BlpR and BlpS monomers is indicated with dark arrowhead. A possible BlpS dimer is indicated by a white arrowhead.

**Sup Fig 4:**
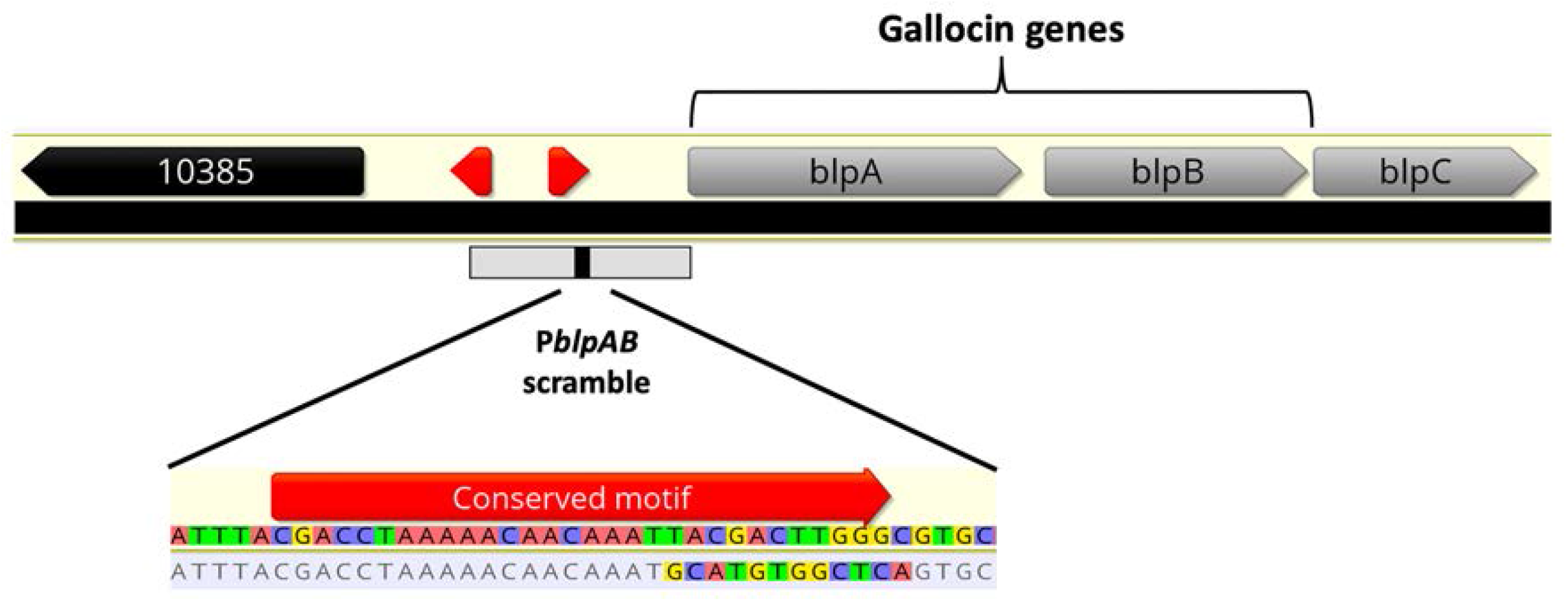
Alignment between P*blpAB* and P*blpAB*-scr. Alignment of the P*blpAB*-scrambled promoter on the WT promoter region around the 30 bp conserved motif.

**Sup Fig 5:**
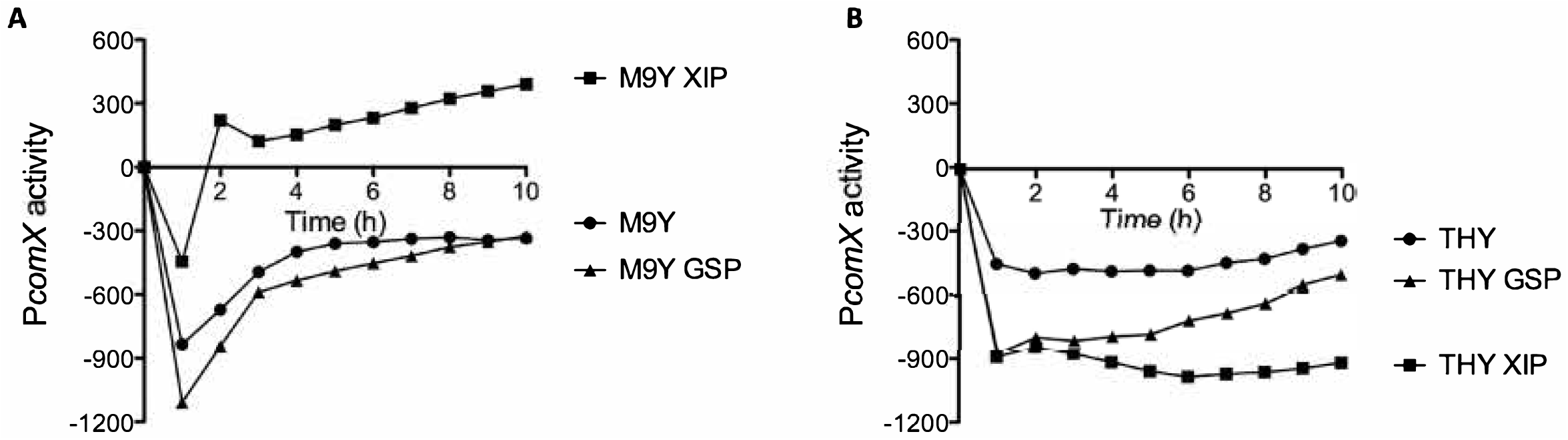
XIP but not GSP activates *comX* promoter. **A**: *comX* promoter (P*comX*) activity in *Sgg* UCN34 pTCV ΩP*comX-gfp* in presence of XIP (1 mM) or GSP (20 nM) in M9Y. **B**: *comX* promoter (P*comX*) activity in *Sgg* UCN34 pTCVΩP*comX-gfp* in presence of XIP (1 mM) or GSP (20 nM) in THY.

## REFERENCES

1. Hoen B, Chirouze C, Cabell CH, Selton-Suty C, Duchêne F, Olaison L, Miro JM, Habib G, Abrutyn E, Eykyn S, Bernard Y, Marco F, Corey GR, International Collaboration on Endocarditis Study Group. 2005. Emergence of endocarditis due to group D streptococci: findings derived from the merged database of the International Collaboration on Endocarditis. Eur J Clin Microbiol Infect Dis Off Publ Eur Soc Clin Microbiol 24:12–16.

2. Boleij A, Tjalsma H. 2012. Gut bacteria in health and disease: a survey on the interface between intestinal microbiology and colorectal cancer. Biol Rev 87:701–730.

3. Corredoira-Sánchez J, García-Garrote F, Rabuñal R, López-Roses L, García-País MJ, Castro E, González-Soler R, Coira A, Pita J, López-Álvarez MJ, Alonso MP, Varela J. 2012. Association Between Bacteremia Due to Streptococcus gallolyticus subsp. gallolyticus (Streptococcus bovis I) and Colorectal Neoplasia: A Case-Control Study. Clin Infect Dis 55:491–496.

4. Aymeric L, Donnadieu F, Mulet C, du Merle L, Nigro G, Saffarian A, Bérard M, Poyart C, Robine S, Regnault B, Trieu-Cuot P, Sansonetti PJ, Dramsi S. 2018. Colorectal cancer specific conditions promote Streptococcus gallolyticus gut colonization. Proc Natl Acad Sci U S A 115:E283–E291.

5. Kommineni S, Bretl DJ, Lam V, Chakraborty R, Hayward M, Simpson P, Cao Y, Bousounis P, Kristich CJ, Salzman NH. 2015. Bacteriocin production augments niche competition by enterococci in the mammalian GI tract. Nature 526:719–722.

6. Heng NCK, Wescombe PA, Burton JP, Jack RW, Tagg JR. 2007. The Diversity of Bacteriocins in Gram-Positive Bacteria, p. 45–92. In Riley, PDMA, Chavan, DMA (eds.), Bacteriocins. Springer Berlin Heidelberg.

7. Oppegård C, Rogne P, Emanuelsen L, Kristiansen PE, Fimland G, Nissen-Meyer J. 2007. The two-peptide class II bacteriocins: structure, production, and mode of action. J Mol Microbiol Biotechnol 13:210–219.

8. de Saizieu A, Gardès C, Flint N, Wagner C, Kamber M, Mitchell TJ, Keck W, Amrein KE, Lange R. 2000. Microarray-Based Identification of a Novel Streptococcus pneumoniae Regulon Controlled by an Autoinduced Peptide. J Bacteriol 182:4696–4703.

9. van der Ploeg JR. 2005. Regulation of Bacteriocin Production in Streptococcus mutans by the Quorum-Sensing System Required for Development of Genetic Competence. J Bacteriol 187:3980–3989.

10. Shanker E, Federle MJ. 2017. Quorum Sensing Regulation of Competence and Bacteriocins in Streptococcus pneumoniae and mutans. Genes 8.

11. Morrison DA, Guédon E, Renault P. 2013. Competence for Natural Genetic Transformation in the Streptococcus bovis Group Streptococci S. infantarius and S. macedonicus. J Bacteriol 195:2612–2620.

12. Harrington A, Tal-Gan Y. 2018. Identification of Streptococcus gallolyticus subsp. gallolyticus (Biotype I) Competence-Stimulating Peptide Pheromone. J Bacteriol 200.

13. Rusniok C, Couvé E, Cunha VD, Gana RE, Zidane N, Bouchier C, Poyart C, Leclercq R, Trieu-Cuot P, Glaser P. 2010. Genome Sequence of Streptococcus gallolyticus: Insights into Its Adaptation to the Bovine Rumen and Its Ability To Cause Endocarditis. J Bacteriol 192:2266–2276.

14. van Belkum MJ, Worobo RW, Stiles ME. 1997. Double-glycine-type leader peptides direct secretion of bacteriocins by ABC transporters: colicin V secretion in Lactococcus lactis. Mol Microbiol 23:1293–1301.

15. Danne C, Dubrac S, Trieu-Cuot P, Dramsi S. 2014. Single cell stochastic regulation of pilus phase variation by an attenuation-like mechanism. PLoS Pathog 10:e1003860.

16. Jans C, Meile L, Lacroix C, Stevens MJA. 2015. Genomics, evolution, and molecular epidemiology of the Streptococcus bovis/Streptococcus equinus complex (SBSEC). Infect Genet Evol J Mol Epidemiol Evol Genet Infect Dis 33:419–436.

17. Nissen-Meyer J, Oppegård C, Rogne P, Haugen HS, Kristiansen PE. 2010. Structure and Mode-of-Action of the Two-Peptide (Class-IIb) Bacteriocins. Probiotics Antimicrob Proteins 2:52–60.

18. Fontaine L, Goffin P, Dubout H, Delplace B, Baulard A, Lecat-Guillet N, Chambellon E, Gardan R, Hols P. 2013. Mechanism of competence activation by the ComRS signalling system in streptococci. Mol Microbiol 87:1113–1132.

19. Nikolskaya AN, Galperin MY. 2002. A novel type of conserved DNA-binding domain in the transcriptional regulators of the AlgR/AgrA/LytR family. Nucleic Acids Res 30:2453–2459.

20. Xie Z, Okinaga T, Niu G, Qi F, Merritt J. 2010. Identification of a Novel Bacteriocin Regulatory System in Streptococcus mutans. Mol Microbiol 78:1431–1447.

21. Zou Z, Qin H, Brenner AE, Raghavan R, Millar JA, Gu Q, Xie Z, Kreth J, Merritt J. 2018. LytTR Regulatory Systems: A potential new class of prokaryotic sensory system. PLOS Genet 14:e1007709.

22. Kjos M, Miller E, Slager J, Lake FB, Gericke O, Roberts IS, Rozen DE, Veening J-W. 2016. Expression of Streptococcus pneumoniae Bacteriocins Is Induced by Antibiotics via Regulatory Interplay with the Competence System. PLoS Pathog 12.

23. Danne C, Guérillot R, Glaser P, Trieu-Cuot P, Dramsi S. 2013. Construction of isogenic mutants in Streptococcus gallolyticus based on the development of new mobilizable vectors. Res Microbiol 164:973–978.

24. Martin M. 2011. Cutadapt removes adapter sequences from high-throughput sequencing reads. 1. EMBnet. journal 17:10–12.

25. Langmead B, Trapnell C, Pop M, Salzberg SL. 2009. Ultrafast and memory-efficient alignment of short DNA sequences to the human genome. Genome Biol 10:R25.

26. Liao Y, Smyth GK, Shi W. 2014. featureCounts: an efficient general purpose program for assigning sequence reads to genomic features. Bioinforma Oxf Engl 30:923–930.

27. Love MI, Huber W, Anders S. 2014. Moderated estimation of fold change and dispersion for RNA-seq data with DESeq2. Genome Biol 15:550.

28. Benjamini Y, Hochberg Y. 1995. Controlling the False Discovery Rate: A Practical and Powerful Approach to Multiple Testing. J R Stat Soc Ser B Methodol 57:289–300.

29. Spriestersbach A, Kubicek J, Schäfer F, Block H, Maertens B. 2015. Chapter One - Purification of His-Tagged Proteins, p. 1–15. In Lorsch, JR (ed.), Methods in Enzymology. Academic Press.

30. Devaux L, Sleiman D, Mazzuoli M-V, Gominet M, Lanotte P, Trieu-Cuot P, Kaminski P-A, Firon A. 2018. Cyclic di-AMP regulation of osmotic homeostasis is essential in Group B Streptococcus. PLoS Genet 14.

31. Poyart C, Trieu-Cuot P. 1997. A broad-host-range mobilizable shuttle vector for the construction of transcriptional fusions to β-galactosidase in Gram-positive bacteria. FEMS Microbiol Lett 156:193–198.

